# Automated Enrichment of Phosphotyrosine Peptides for High-Throughput Proteomics

**DOI:** 10.1101/2023.01.05.522335

**Authors:** Alexis Chang, Mario Leutert, Ricard A. Rodriguez-Mias, Judit Villén

## Abstract

Phosphotyrosine (pY) enrichment is critical for expanding fundamental and clinical understanding of cellular signaling by mass spectrometry-based proteomics. However, current pY enrichment methods exhibit a high cost per sample and limited reproducibility due to expensive affinity reagents and manual processing. We present rapid-robotic phosphotyrosine proteomics (R2-pY), which uses a magnetic particle processor and pY superbinders or antibodies. R2-pY handles 96 samples in parallel, requires 2 days to go from cell lysate to mass spectrometry injections, and results in global proteomic, phosphoproteomic and tyrosine specific phosphoproteomic samples. We benchmark the method on HeLa cells stimulated with pervanadate and serum and report over 4000 unique pY sites from 1 mg of peptide input, strong reproducibility between replicates, and phosphopeptide enrichment efficiencies above 99%. R2-pY extends our previously reported R2-P2 proteomic and global phosphoproteomic sample preparation framework, opening the door to large-scale studies of pY signaling in concert with global proteome and phosphoproteome profiling.

## Introduction

Protein phosphorylation plays critical roles in the regulation of many cellular processes and is central to cellular signaling^1–5^. In particular, tyrosine phosphorylation plays a critical role in transduction of extracellular signals to elicit transcriptional changes that drive cellular growth, differentiation, and apoptosis^4,5^. Accordingly, aberrant kinase activity drives many cancers^6,7^ and tyrosine kinases are over-represented in a census of cancer genes^6^. Despite the central role of tyrosine phosphorylation in initiating and maintaining critical cellular processes, phosphotyrosine is low in abundance in comparison to phosphoserine and phosphothreonine due to a lower number of sites and rapid removal by protein tyrosine phosphatases^8,9^.

The lower relative abundance of phosphotyrosine results in relatively few identifications of phosphotyrosine containing peptides (<1%) from global phosphopeptide enrichment^8^. Global phosphopeptide enrichment typically relies on the general affinity of immobilized metal or metal oxide interactions for negatively charged phosphate groups. Higher abundance phosphoserine and phosphothreonine peptides outcompete phosphotyrosine peptides during both enrichment and detection by mass spectrometry^8^. As a result, study of phosphotyrosine signaling is more successful when phosphotyrosine-specific enrichment is used^8^. Methods for phosphotyrosine enrichment have traditionally relied on phosphotyrosine-specific antibodies. While effective, phosphotyrosine specific antibodies are expensive, limiting feasibility of larger scale studies. An alternative to phosphotyrosine-specific antibodies are SH2 protein domains which naturally bind phosphotyrosine at low affinity^10–13^ and have been engineered to increase affinity for use as reagents. Engineered SH2 domains have been shown to enrich phosphotyrosine peptides comparable or better than antibodies^13–20^. Such engineered SH2 domains are termed phosphotyrosine superbinder SH2 or sSH2 domains. sSH2 domains can be recombinantly expressed and purified from *Escherichia coli*, reducing their production cost relative to antibodies.

While phosphotyrosine enrichment methods have been developed and widely used, most methods are manual^8,14,15,17,21–27^ and/or low throughput^16^, limiting cost-effectiveness, scalability and reproducibility. For similar reasons, proteomic sample preparation and phosphopeptide enrichments have been fully automated in high throughput formats^28–30^. Development of a fully automated phosphotyrosine enrichment method with cost effective affinity reagents will help expand studies of phosphotyrosine signaling. Recently, Martyn *et al*. (2022) automated a workflow for phosphotyrosine enrichment using engineered SH2 domains bound to magnetic beads on a robotic magnetic particle processor (Thermo Fisher KingFisher Flex™)^20^. However, as most commonly used proteomic sample preparation workflows, this method still relies on some manual steps for sample clean-up. Other groups have established different combinations of global phosphopeptide enrichment and pY-specific affinity reagents for clean up of phosphotyrosine peptide samples^14–16,19,23^. However, sequential phosphotyrosine and global phosphopeptide enrichment steps remain to be implemented in a fully automated workflow. Here we develop a scalable, reproducible, and automated procedure for efficient enrichment and clean up of phosphotyrosine containing peptides and show full integration with automated proteomic and global phosphoproteomic sample preparation.

We developed and benchmaked an automated and high throughput phosphotyrosine enrichment module called rapid-robotic phosphotyrosine proteomics (R2-pY) that utilizes the Src SH2 superbinder domain (sSrc) or anti-phosphotyrosine antibodies in combination with a robotic magnetic bead handler capable of processing 96 samples in parallel. In addition to reducing hands on processing with automation, the current implementation of R2-pY reduces affinity reagent cost five fold, with further cost reduction possible. Additionally, R2-pY integrates with automated proteomic and phosphoproteomic sample preparation such that systematic measurement of protein abundance and phospho signaling is achieved at the end of the workflow. We benchmark R2-pY with pervanadate and serum stimulated HeLa cells. We highlight that R2-pY is quantitatively scalable to allow flexibility in terms of sample input amounts to a range of users. We also highlight the high reproducibility, enrichment selectivity and cost efficiency of R2-pY. We provide detailed methods for implementation including integration within the previously published R2-P1 and R2-P2 workflows for efficient measurement of cellular signaling across many samples.

## Methods

### Preparation of superbinders

Sequences encoding the human Src and Grb2 SH2 superbinder domains and tandem construct were integrated in the pET28a(+) vector with an N-terminal His6 tag and thrombin cleavage site (sequences deposited in Supporting File 1). The Src SH2 superbinder (sSrc) contains mutations T8V, C10A, and K15L^13^. The Grb2 SH2 superbinder (sGrb2) contains mutations A8V, S10A, and K15L^13^. The tandem construct consists of the Src SH2 domain followed by the Grb2 SH2 domain (Supporting File 1). Each superbinder SH2 construct (sSH2) was expressed in T7 Express Competent E. *coli* cells (New England Biolabs). Protein expression was induced at OD_600 nm_ 0.6 - 0.8 with 1 mM IPTG overnight at 18°C. Cell pellets were harvested by centrifugation and stored at - 80°C until purification. Cell pellets were resuspended in a lysis buffer containing phosphate-buffered saline (PBS) (0.01 M phosphate buffer, 0.0027 M potassium chloride, 0.137 M sodium chloride, pH 7.4), 0.1 mg/mL lysozyme, and 10 U/mL benzonase nuclease. Cell suspensions were incubated for 10 min at 4°C, sonicated on ice at 40% power using six 30-sec pulses with 1-min intervening rests then incubated for 10 min further at 4°C. Lysate was clarified by centrifugation at 7197 x *g* for 60 min and 0.45 μm filtration. sSH2 domains were immediately purified by the N-terminal His6 tag with TALON^®^ resin (Takara Bio) using a gravity column (detailed purification method in the Supporting Methods 1). Briefly TALON resin was equilibrated in 1X PBS, pH 7.4. Approximately 2 mL of dry resin volume was used per 1 L culture or 5 g of cell pellet. Lysate was incubated with resin for 30 - 60 min at 4 - 8°C with gentle mixing. Flow through was collected then the column was washed sequentially with 1X PBS pH 7.4, 1X PBS pH 7.4 with additional 150 mM NaCl, 1X PBS, 5 mM imidazole, pH 7.4, optional 1X PBS, 10 mM imidazole, pH 7.4, and again with 1X PBS, pH 7.4. Following washes, sSrc was eluted in six to eight column volumes of 1X PBS, 300 mM imidazole, pH 7.4. Elution fractions were pooled based on sodium dodecyl sulfate polyacrylamide gel electrophoresis (SDS PAGE) and buffer exchanged into PBS, 0.05% sodium azide, pH 7.4 then concentrated to 2 mg/mL with a 3 kDa centrifugal filter. Protein concentration was determined by absorbance at 280 nm on a NanoDrop^TM^ spectrophotometer with a mass extinction coefficient of 0.981 for Src sSH2, 0.980 for Grb2 sSH2, and 1.06 for tandem sSH2. A representative SDS PAGE of the clarified lysate and purified Src sSH2 along with purified fractions of Grb2 sSH2 and tandem sSH2 is included for reference (Supporting Figure 1A and Supporting Figure 1B). Purified constructs were aliquoted and stored at −80°C until conjugation. Prior to conjugation, sSH2s were buffer exchanged into 50 mM borate pH 8.5 and concentrated to 1.75 to 2 mg/mL with a 3 kDa centrifugal concentrator at 4°C. sSH2 constructs were immobilized on NHS-Activated Magnetic beads (Pierce™/ Thermo Scientific) in the following way. NHS-activated magnetic bead slurry (10 mg/mL) and purified sSH2 constructs were equilibrated to room temperature for 30 minutes. Bead slurry aliquots of 300 μL were transferred to microtubes. Storage solution was removed. 1 mL of ice-cold hydrochloric acid was added and vortexed gently with beads for 15 seconds. Hydrochloric wash was discarded followed immediately by addition of 300 μL of purified sSH2 solution (purified sSH2 at 1.5-2 mg/mL in 50 mM borate pH 8.5). Beads were resuspended by gentle flicking, inversion and vortexing for 5 - 15 sec. Beads and proteins incubated with rotation for 2 hours at room temperature. Protein supernatant was saved for assessment of conjugation efficiency by SDS PAGE and 660 nm assay. Beads were quenched by three sequential washes with 50 mM Tris, 100 mM NaCl, pH 8.0; each wash was incubated with beads for 5 min, 30 min and 85 min, respectively at room temperature with inversion. Conjugated beads were resuspended at dilute condition (3 to 4 mg/mL bead slurry) with 1 mL of 50 mM Tris, 100 mM NaCl, pH 8.0 and stored at 4°C until R2-pY.

### Quantification of conjugation efficiency

Quantification of conjugation efficiency was performed from saved aliquots of load and flow through with 660 nm reagent (Pierce™). A batch of NHS-conjugated sSrc magnetic beads was prepared as described above and conjugation efficiency measured as 147.8 μg of sSrc (or 10 nmol sSrc) per mg of dry beads. This equates to 0.1 nmol of sSrc per μL of bead slurry.

The Pierce™ 660 nm assay was performed in a standard 96-well microplate as recommended^31^. Briefly, the standard curve was prepared with TALON^®^-purified sSrc in 1X PBS pH 7.3 at the following concentrations (mg/mL): 2, 1.5, 1, 0.75, 0.5, 0.25, 0.125, 0.025. Saved aliquots of sSrc from before and after incubation with Pierce™ NHS-activated magnetic beads were added at 10 μL per well in duplicate. Two-fold and fourfold dilutions of sSrc at 2 mg/mL from before incubation were made in 50 mM borate pH 8.5 to stay within the linear assay range. Then 150 μL of 660 nm protein assay reagent was added to each well and the plate was mixed at medium speed for 1 min followed by 5 min without mixing at room temperature. Absorbances were measured at 660 nm in triplicate then averaged. A four-parameter standard curve was fitted to determine the protein concentration of each unknown sample.

### Cell culture and treatment

HeLa S3 cells were cultured at 37°C and 5% CO2 in Dulbecco’s modified Eagle’s medium (DMEM) supplemented with 4.5 g/L glucose, L-glutamine, and 10% fetal bovine serum (FBS) and 0.5% streptomycin / penicillin. To generate bulk phosphopeptides for method comparisons, cells were grown to 80% confluency, incubated in serum-free medium for 6 hours prior to treatment with 1 mM pervanadate for 15 min, followed by addition of 10% FBS for 15 min. At the time of harvest, cells were rinsed three times quickly with ice-cold PBS, resuspended in minimal volume of PBS, transferred to conical tubes, centrifuged at 2000 x *g* for 5 min at 4°C. PBS wash was discarded and cells were flash frozen in liquid nitrogen prior to storage at −80°C.

### Cell lysis, protein reduction and alkylation

Cell lysis was performed in 8 M urea, 50 mM HEPES, 75 mM NaCl, pH 8.0. Cell resuspension was sonicated with 6 x 20 sec pulses at 5-6 watts with 20-30 sec rests, all while submerged in ice. Lysate was clarified by centrifugation at 7197 x *g* for 25 min at 20°C. Protein content was assayed using the bicinchoninic acid method (Pierce™). Proteins were reduced with 5 mM dithiothreitol (DTT) for 30 min at 55°C, alkylated with 15 mM iodoacetamide for 15 min at room temperature in the dark, then quenched with 5 mM DTT for 15 min at room temperature.

### Peptide digestion and desalting

Digestion of proteins to peptides was performed either in bulk solution followed by desalting on SEP-PAK C18 cartridges (Waters^TM^) or on-beads following an optimized R2-P1 protocol in 96-well plates on the KingFisher^TM^ Flex robot (optimized R2-P1 protocol in the Supporting Methods 1)^28^ (Figure 1). In-solution digestion and cartridge desalting were employed in procedural optimization experiments, specifically for testing impact of: different conjugation chemistries (Figure 2), different superbinder constructs (Figure 3), different sSrc amounts (Figure 4) and different peptide input amounts for tests of quantitative scalability (Figure 5). Otherwise, R2-P1 was employed for peptide digestion and desalting (Figure 6).

When peptides were digested in-solution, the reduced and alkylated lysate was diluted to reach less than 1M urea in 50 mM ammonium bicarbonate pH 8.0. Trypsin was added to reach a final trypsin:protein ratio of 1:100 by mass and a final trypsin concentration greater than 5 ng/mL. Digests were mixed briefly by inversion and pH was measured to ensure pH between 8 and 8.5. Digests were incubated overnight (15 hours) at 37°C in conical tubes without constant agitation. Digestion was quenched by addition of 10% TFA to reach a final concentration of 0.5% TFA for a pH < 2. Quenched digests were centrifuged at 7,197 x *g* for 11 min at room temperature to remove precipitates. Peptides were desalted on Waters^TM^ SEP-PAK C18 cartridges in the following way. Columns were activated by sequential addition of 1 column volume (CV) of methanol; 3 CVs of 100% ACN; 1 CV of 70% ACN, 0.25% acetic acid; 1 CV of 40% ACN; 3 CV 0.1% TFA. Acidified digests were then loaded followed by reload of the flow through. Column was washed with 3 CV of 0.1 % TFA. Peptides were eluted with 0.5 CV of 40% ACN, 0.5% AA followed by 0.5 CV of 70% ACN, 0.25% AA. Elutions were pooled, mixed thoroughly then aliquoted into microtubes in either 0.25 mg, 0.5 mg or 1 mg fractions ahead of vacuum centrifugation. Dried peptides were stored at −20°C until further enrichment.

When R2-P1 was used, protein extracts were diluted to 5 mg/mL in HeLa cell lysis buffer (8 M urea, 50 mM HEPES, 75 mM NaCl, pH 8.0) and centrifuged to remove precipitates. The R2-P1 total proteomic sample preparation workflow^28^ was implemented on the KingFisher^TM^ in the following way:

The 96-well comb was stored in plate #1, lysates, beads and ethanol mixture in plate #2, ethanol washes in plates #3 to #6, elution/digestion buffer in plate #7, and second elution (water) in plate #8. Final ethanol concentration was 80% (v/v) in plates #2 to #6. Each well of plate #2 received 1 mg of pervanadate and serum stimulated HeLa lysate. The method was configured to bind proteins to beads in plate #2 and clean proteins in plates #3 to #6. The protocol paused before protein digestion. Digestion solution was prepared immediately before use at a final volume of 500 μL per well. The automated protocol resumed and beads were moved to plate #7 for elution/digestion at 37°C for 3.5 hours with constant agitation. Beads were moved to the second elution in plate #8 and afterward discarded. Upon program end, eluates from plates #7 and #8 were combined and quenched with 60 μL of 10% formic acid, 75% ACN and clarified by centrifugation. At this point small aliquots (i.e. 50 μg) can be saved for total proteome measurement. We note total proteome samples particularly benefit from ensuring pH < 2. Total proteome samples were filtered through C8 or C18 stage tips to avoid carryover of magnetic beads to the LC system^28^. Otherwise, acidified eluates were dried by vacuum centrifugation ahead of pY-peptide enrichment with R2-pY.

### R2-pY phosphotyrosine peptide enrichment

The R2-pY phosphotyrosine enrichment was implemented on the KingFisher™ Flex in the following way: The 96-well comb and pY capture beads (antibody or sSrc conjugated magnetic beads) were stored in plate #1, resuspended peptides in plate #2, Affinity Purification buffer (1X AP: 20 mM MOPS-NaOH, 10 mM Na_2_HPO_4_, NaCl, pH 7.2) washes in plates #3 to #5, water wash in plate #6, acidic elution-1 in plate #7 and acidic/organic elution-2 in plate #8 (Figure 1). Shallow 96-well KingFisher plates were used for plates #1, #7 and #8 and deep 96-well plates were used for plates, #2 to #6. p-Tyr-100 and p-Tyr-1000 antibodies were purchased as magnetic bead conjugates from Cell Signaling Technology^®^ and mixed in a 1:1 ratio by volume, washed in 1X AP and used at a ratio of 83 μL bead slurry per 1 mg of peptides. Amounts of sSrc beads used per well were varied during the optimization experiments and settled on 83 μL of 10mg/mL bead slurry corresponding to approximately 8.3 nmol of conjugated sSrc protein when conjugation efficiency measures 0.1 nmol of sSrc per μL of bead slurry. The method was configured to collect the magnetic beads from plate #1, move them to plate #2 for pY-peptide binding, then to plates #3-6 for washing, and subsequently to plate #7 for the first elution. The pY-binding, washing and first elution take 140 min, then the protocol pauses to allow for loading of plate #8 which contains the acidic/organic elution-2. Elution-2 is prepared immediately before use to minimize evaporation of acetonitrile. The beads are then moved to plate #8 for a second elution that takes 11.3 min. Plates are removed from the robot at this point and the elution 1 and 2 are pooled and brought to 80% ACN followed by 10 min centrifugation at >16,000 x *g* and 25°C. Pooled, adjusted, and clarified R2-pY-eluates are immediately subjected to R2-P2 with Fe^3+^ magnetic beads.

**Figure 1.**
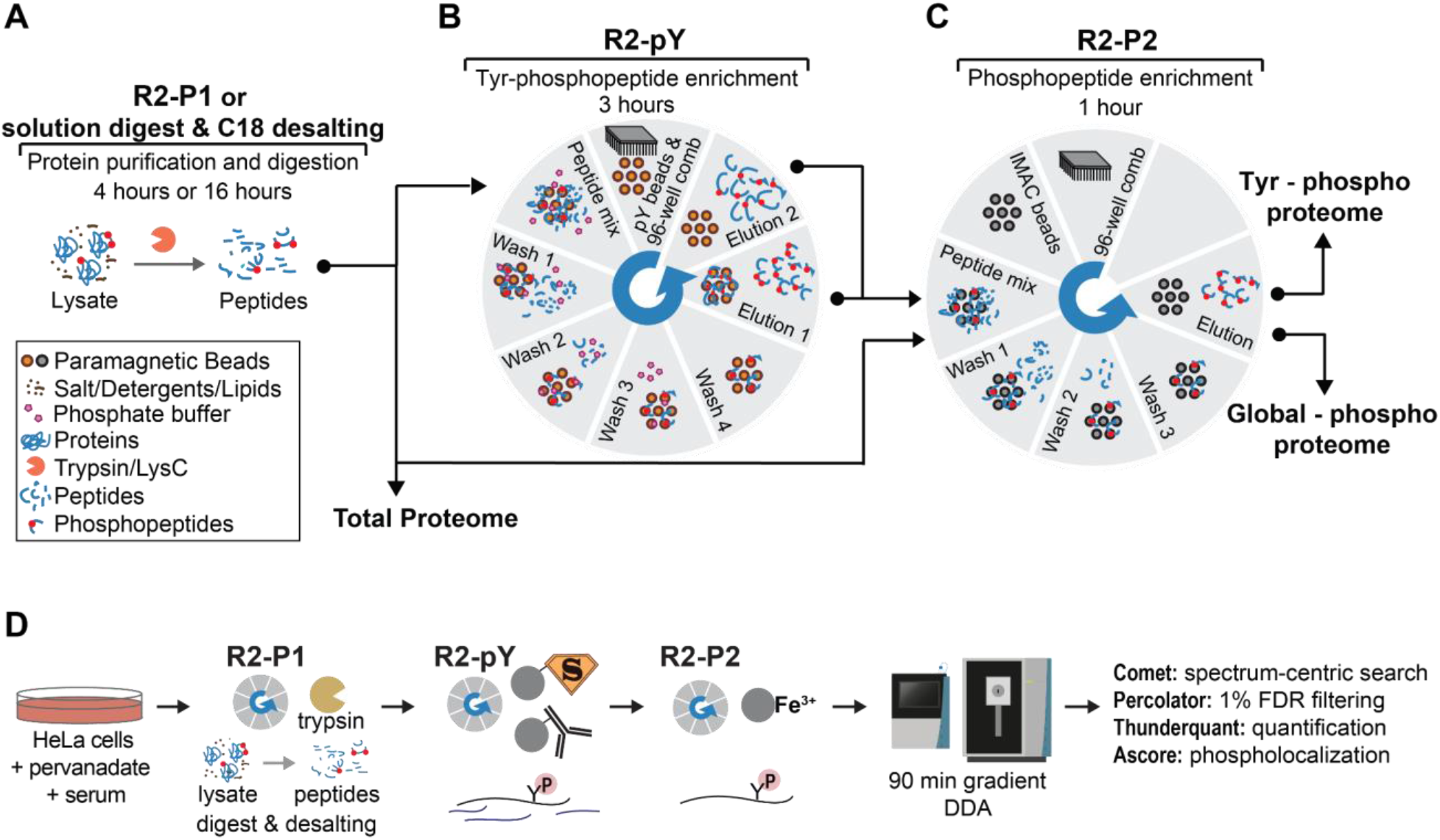
Overview of R2-pY module. R2-P1, R2-pY and R2-P2 provide a unified platform for preparation of each total proteome, global phosphoproteome and tyrosine phosphoproteome samples. **A.** Proteins from the sample are digested either with the R2-P1 module or in-solution digestion and cartridge desalting. **B**. Configuration of R2-pY. Desalted peptides are resuspended in an affinity purification buffer then incubated with either sSrc or anti-pY antibody magnetic beads. Then, bead-bound peptides are washed before elution in the last two plates. **C**. Configuration of the R2-P2 module^28^. R2-P2 serves dual purposes: Within pY enrichment, the pooled elution from R2-pY is added to plate 3 of R2-P2 for further enrichment. Alternatively, addition of desalted total proteome peptides to R2-P2 generates global phosphoproteome samples. **D.** Workflow for validation of the new R2-pY module.

### R2-P2 clean-up of R2-pY eluate

Cleanup with R2-P2 was performed on the KingFisher™ Flex in the following way: The 96-well comb is stored in plate #1, magnetic Fe^3+^-IMAC beads in plate #2, adjusted R2-pY eluate in plate #3, wash solutions in plates #4 to #6, and elution solution in plate #7 (Figure 1). Shallow 96-well KingFisher plates were used for all plates except plate #3 which used a deep 96-well. The method was configured to collect the magnetic beads in plate #2, move them to plate #3 for phosphopeptide binding and subsequently to plate #4, #5, and #6 for washing. The phosphopeptide purification takes 40 min, and the protocol pauses to allow for loading of plate #7 containing the elution solution, which is prepared immediately before use, to avoid evaporation of ammonia. The beads are then moved to plate #7 where phosphopeptides elute. Plates are removed from the robot at this point, and the user immediately neutralizes the elution by acidification. Eluates were filtered through C8 or C18 stage tips to avoid carryover of magnetic beads to the LC system. Samples were dried by vacuum centrifugation and stored at −20°C until resuspension in 4% formic acid and 3% ACN for LC-MS/MS analysis.

### Conjugation of SH2 domains to magnetic beads

Three different magnetic particle functionalizations were tested for conjugation of sSrc to magnetic particles: NHS, epoxy and maleimide (Supporting Methods 2).

#### NHS conjugation

Conjugation of TALON^®^-purified sSrc to NHS-activated Pierce™ magnetic beads was performed in the following way: sSrc in 50 mM borate pH 8.5 and bead solution were equilibrated to room temperature for 30 min. Beads were activated by brief wash in ice-cold 1 mM hydrochloric acid followed by incubation with sSrc for 2 hours. Following incubation, beads were washed three times with 50 mM Tris, 100 mM NaCl, pH 8.0 within a 2 hour duration. Conjugated beads were stored at 4°C in 50 mM Tris, 100 mM NaCl, pH 8.0 at less than 3 to 4 mg/mL bead slurry. Immediately prior to R2-pY, beads were washed three times in 1X AP pH 7.2 then resuspended in 1X AP pH 7.2 at 10 mg/mL bead slurry.

#### Epoxy conjugation

sSrc was conjugated to PureCube Epoxy-Activated MagBeads (Cube Biotech). TALON^®^-purified sSrc was equilibrated into Epoxy coupling buffer I or II (I: 150 mM NaH_2_PO_4_, 100 mM NaCl, pH 7.2; or, II: 150 mM NaCO_3_, 500 mM NaCl, pH 8.3, respectively) then concentrated to 2 mg/mL. Aliquots of epoxy-activated magnetic beads were prepared by four washes in water followed by one wash in each coupling buffer (I or II). Then, each 1 mL aliquot of 25% bead slurry was resuspended in 0.5 mL of respective coupling buffer I or II. Beads were incubated at room temperature for 10 minutes before addition of 1 mL sSrc (concentrated to 2 mg/mL with coupling buffer I or II; ± TCEP). sSrc was incubated with beads for 24 hours at room temperature with end-over-end mixing. Beads were quenched with 0.5 mL of 1M Tris, pH 7.4 for 10 hours at room temperature with end-over-end mixing. Beads were washed five times with respective coupling buffers and resuspended in 0.5 mL storage solution (20 mM sodium acetate, 130 mM sodium chloride, pH 6.5), pooled, and stored at 4°C until R2-pY.

#### Maleimide conjugation

sSrc was conjugated to PureCube Maleimide-Activated MagBeads (Cube Biotech) in the following way: sSrc was equilibrated into 100 mM sodium acetate, 150 mM NaCl, pH 6.5 and concentrated to 2 mg/mL. Then 2 mL of sSrc at 2 mg/mL was added to 4 mL of 25% bead slurry to yield 4 mg of sSrc per 1 mL dry bead volume. Protein and beads were mixed end-over-end at room temperature for 2 hours. Beads were then washed three times with 100 mM sodium acetate, 150 mM NaCl, 10 mM cysteine, pH 5.3 in which the second wash was incubated for 30 min at room temperature with mixing. Beads were washed three times with 50 mM borate, 100 mM NaCl, pH 8.5 then resuspended in 100 mM sodium acetate, 150 mM NaCl, pH 6.5 at 25% to 40% bead slurry and stored at 4°C until R2-pY. Beads were resuspended to 25% slurry ahead of aliquoting for R2-pY.

### Mass spectrometry acquisition

Dried peptide and phosphopeptide samples were dissolved in 4% formic acid, 3% acetonitrile and analyzed on Orbitrap Eclipse (Thermo Fisher) or Orbitrap Exploris 480 Mass Spectrometer (Thermo Fisher) equipped with an Easy-nLC 1200 system. Only samples comparing conjugation chemistries were measured on the Orbitrap Exploris 480 Mass Spectrometer. Peptides were loaded onto a 100 μm ID x 3 cm precolumn packed with Reprosil C18 3 μm beads (Dr. Maisch GmbH), and separated by reverse-phase chromatography on a 100 μm ID x 30 cm analytical column packed with Reprosil C18 1.9 μm beads (Dr. Maisch GmbH) and housed into a column heater set at 50°C. Phosphopeptides were separated by a 90 minute gradient where percentage acetonitrile in 0.125% formic acid ranged from 3.2% to 22.4% for 71 minutes, 22.4% to 48% for 2 minutes, 76% for 5 min and 2.4% for 10 min.

Full MS scans were acquired from 375 to 1500 m/z at 120,000 resolution. The fill target was set to ‘Standard’. The maximum injection time was set to ‘Auto’. The maximum number of ions on the full MS scan were selected for fragmentation within a 3 sec cycle time using 1.6 m/z precursor isolation windows and beam-type collisional-activation dissociation (HCD) with 30% normalized collision energy. MS/MS spectra were collected at 50,000 resolution with fill target set to 1e5 ions and maximum injection time of 86 ms. Fragmented precursors were dynamically excluded from selection for 45 s. Data was stored in centroid mode.

### Mass spectrometry data analysis

DDA-MS/MS spectra were searched with Comet (2019.01.rev.2^32^) against the human proteome. The human proteome sequence database was retrieved from Uniprot in 2014 and contains 87,613 entries which include reviewed, unreviewed and isoform sequences. Peptides were searched with trypsin digestion allowing up to 4 missed cleavages. The precursor m/z mass tolerance was set to 50 ppm except for comparisons of conjugation chemistry which was searched with 20 ppm precursor tolerance. Fragment m/z mass tolerance was set to 0.02 Da. Constant modification of cysteine carbamidomethylation (57.02146372118 Da) and variable modification of n-term acetylation (42.01056468472 Da), methionine oxidation (15.9949146202 Da), and phosphorylation of serine, threonine and tyrosine (79.96633089136 Da) were used for all searches. Search results were filtered to a 1% FDR at PSM level using Percolator^33^. Phosphorylation sites were localized using an in-house implementation of the Ascore algorithm^34^. Phosphorylation sites with an Ascore > 13 (*P* < 0.05) were considered confidently localized. Peptides were quantified using in-house software measuring MS1 chromatographic peak maximum intensities.

### Bioinformatic analysis

Bioinformatic analysis was performed using R (https://r-project.org/). Quantitative values obtained from DDA analyses were derived from maximum intensity PSMs for the most prevalent charge states per peptide and were sample median normalized before correlation and variation analyses.

For all boxplots, the lower and upper hinges of the boxes correspond to the 25th and 75th percentiles, and the bar in the box corresponds to the median. The upper and lower whiskers extend from the highest and lowest values, respectively, but no further than 1.5 times the IQR from the hinge. All correlation calculations utilize Pearson’s method.

Analysis of sequence preference of pY affinity reagents was performed with IceLogo^35^ with a p-value cutoff of 0.05. The sets of pY peptides enriched by each sSrc and antibodies were mapped back to the full protein and extended 6 amino acids on both sides of pY sites prior to enrichment analysis against a human proteome FASTA file. Only pY-peptides with at least one confidently localized pY modification were used in the analysis. Sites identified in multiple replicates were multiply counted in a set to reflect redundancy in sequence preference. The human proteome background set was sampled regionally with phosphotyrosine at position 7 as the anchor. Within the resulting logos, the differences in amino acid frequency between the pool of enriched pY peptides and the human proteome are represented by the individual amino acid heights.

Representation of pY sites detected across the human kinome was visualized with Coral using the web interface: phanstiel-lab.med.unc.edu/Coral ^36^. The set of unique peptides with at least one confidently localized phosphotyrosine site were used in the analysis. The human kinase associated with each pY-peptide served as Coral input. Kinases represented by pY-peptides enriched by either sSrc or antibodies were combined for representation in a single plot.

## Results and discussion

### Overview of the R2-pY method

R2-pY provides an automated workflow for enrichment of phosphotyrosine-modified peptides. We previously developed a high-throughput and automated method using magnetic particles for proteomic and global phosphoproteomic sample preparation^28^. Here we further expand the PTM enrichment capabilities of this workflow, by developing an automated phosphotyrosine (pY) peptide enrichment module. Along with the proteomic and global phosphoproteomic preparation methods, the phosphotyrosine enrichment module utilizes a magnetic particle processing robot with capacity for 96 samples. The procedure offers the flexibility to use commercially available antibodies conjugated to magnetic beads^37,38^ or SH2 domain-derived pY superbinders, which we conjugate to NHS-activated magnetic beads.

The pY enrichment module starts from desalted peptides, enriches pY-modified peptides, and then performs immobilized metal affinity chromatography (IMAC) to further remove non-phosphorylated peptides and cleanup salts prior to mass spectrometry acquisition. We designed the R2-pY workflow to allow protein extract samples to be processed by R2-P1^28^ or by in-solution-digestion and solid-phase extraction desalting (Figure 1A). Desalted peptides are dried and resuspended in an affinity purification (AP) buffer. The R2-pY protocol starts by incubating pY-capture beads with peptides (Figure 1B). The bead-peptide complexes are then passed through three plates of AP buffer to wash off peptides with no pY modifications (Figure 1B). A fourth wash with water removes the AP buffer prior to pY-peptide elution (Figure 1B). pY-peptides are eluted from beads with 0.5% to 1 % trifluoroacetic acid (TFA) (Figure 1B). Eluates are then processed by R2-P2^28^ using Fe^3+^-IMAC beads for stringent removal of non-phosphorylated peptides and residual AP reagent (Figure 1C). Overall the R2-pY to R2-P2 workflow takes 4 hours and results in peptides that can be dried and analyzed by LC-MS/MS. Detailed protocols for R2-pY and R2-P2 and KingFisher™ Flex programs are provided in the methods and Supporting Methods 1.

In the following sections we present results from a series of parameter variations that both inform development and flexibility of R2-pY. Each parameter optimization experiment was tested with a standardized tryptic digest of whole cell HeLa lysate in which HeLa cells were treated with pervanadate and serum. Pervanadate inhibits protein tyrosine phosphatases providing a sample with high phosphotyrosine content to facilitate method benchmarking^8,15,23^. In the end we demonstrate performance of the fully automated robotic pY-peptide preparation workflow consisting of R2-P1 to R2-pY to R2-P2, which starts from cell lysate and results in MS-ready pY-peptide samples (Figure 1D).

### Preparation of pY superbinders in E. *coli*

To obtain pure and soluble pY superbinders we designed superbinder expression constructs based on previous enrichment methods that utilized pY superbinders^14,15,17^. We optimized conditions for soluble expression and purified superbinders from E. *coli* lysate via an N-terminal His6 tag (Figure 2A). Src superbinder (sSrc) seemed to express well in the soluble fraction and we obtained around 20 mg of purified sSrc from 1 L of E. *coli*. After buffer exchange, pY superbinders can be flash frozen and stored at −80°C for at least several months before conjugation to magnetic beads (Figure 2A).

**Figure 2.**
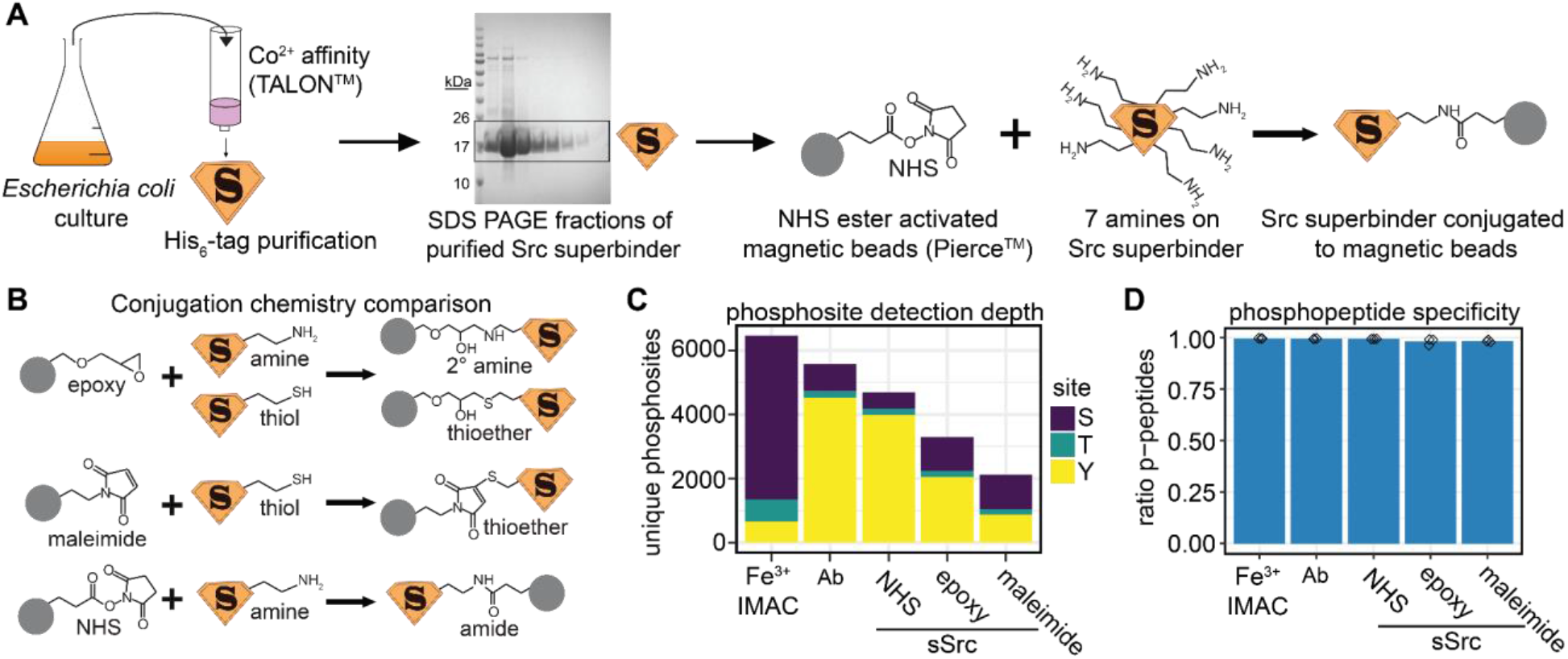
Conjugation chemistry and Fe^3+^-IMAC clean-up impact enrichment success. **A.** Workflow for preparation of superbinder and conjugation to magnetic beads. sSrc was expressed in E. *coli* then purified by the His6 tag with cobalt(II) immobilized metal affinity chromatography. Fractions containing pure sSrc were pooled and buffer exchanged then conjugated to Pierce™ NHS-activated magnetic beads. **B.** Three different chemistries were tested for conjugation of sSrc to magnetic beads. **C.** Number of unique phosphosites confidently localized after triplicate enrichments from 1 mg of peptide input with each of the five different capture reagents (n=3). Bead types (*left to right):* Fe^3+^ IMAC represents global phosphopeptide enrichment, pTyr antibody (Ab), and sSrc conjugated with three different chemistries described in (B). **D.** Barplot depicting the ratio of the number of phosphopeptides out of all unique peptides enriched in each sample. Diamonds represent individual replicates (n=3).

### Conjugation chemistry and Fe^3+^-IMAC clean-up impact enrichment success

Previous methods for covalent conjugation of sSrc to beads use cyanogen bromide-activated Sepharose^15,17,18^. For compatibility with R2-P2, we sought to conjugate sSrc to magnetic beads. We evaluated three different bead functionalizations targeting amines (NHS and epoxy) or thiols (maleimide) given sSrc contains 7 lysines and 2 cysteines (Figure 2B). sSrc beads were tested alongside a 1:1 mixture of p-Tyr-100 and p-Tyr-1000 beads for pY-peptide enrichment from 1 mg of peptide input in triplicate. Capture reagents were compared in terms of number of phosphosites (depth) and percentage of peptides that are phosphopeptides (selectivity) (Figure 2C, 2D).

To highlight the improvement in pY selectivity and depth achieved by R2-pY, we compared R2-pY to global phosphopeptide enrichment with Fe^3+^-IMAC using R2-P2 (Figure 2C, 2D). Fe^3+^-IMAC enrichment alone resulted in the identification of 654 unique pY sites whereas both Ab and NHS-conjugated sSrc captured 4528 and 3979 pY sites, respectively (Figure 2C). The epoxy-based and maleimide-based sSrc conjugations captured notably less pY sites (2051 and 878 sites, respectively) (Figure 2C). Lower pY capture efficiency observed by the epoxy and maleimide chemistries may stem from lower conjugation efficiencies. Regardless, all samples displayed exceptionally high phosphopeptide purity (>98% on average) (Figure 2D), likely achieved at the R2-P2 global phosphopeptide clean-up step^8,14,15,19,22–24^. Taken together, both pY antibodies and NHS-conjugated sSrc performed exceptionally well in R2-pY and the combination of R2-pY with R2-P2 delivers high purity phosphopeptides. Based on these results, in subsequent experiments we use the NHS-activated magnetic beads for conjugation of sSH2 domains and perform R2-P2 clean-up on eluates from R2-pY.

### R2-pY allows utilization of different pY capture reagents

After establishing that sSrc beads conjugated via NHS chemistry efficiently enriches pY-peptides, we sought to evaluate alternative superbinder constructs for R2-pY phosphotyrosine peptide enrichment. We compared Src SH2 superbinder (sSrc), Grb2 SH2 superbinder (sGrb2)^13^, and a novel tandem construct we designed consisting of sSrc and sGrb2 connected sequentially (sequences deposited in Supporting File 1). To measure the capture efficiency and sequence bias of each binder, we performed R2-pY to R2-P2 with all three binders alongside the p-Tyr-100 and p-Tyr-1000 antibody mixture. Enrichments were performed in triplicate, each from 1 mg of peptide input.

The different binders displayed distinct efficiency in capturing pY-peptides. Notably, sSrc and tandem yielded high numbers of confidently localized pY sites (4856 and 4491, respectively), which were higher than antibody (3787 pY sites) (Figure 3A), and much higher than sGrb2 (675 pY sites) (Figure 3A). As expected, numbers of unique pY sites correlated with median log2 intensity of phosphopeptides (Figure 3B). sSrc captured not only the highest number of pY sites but also was able to recover the highest quantities of individual pY-peptides. Tandem and antibody still captured both a broad set and high quantities of each unique pY-peptide on average. The differences in numbers of unique enriched pY sites can be explained by differences in pY binding efficiency rather than differences in sample cleanliness as all four binders displayed high phosphopeptide count and intensity-based enrichment: >99.9% for sSrc, tandem and antibody and >95% for sGrb2 (Supporting Figure 2A and 2B). Thus, sSrc remains the most effective sSH2 domain among the three tested and additionally shows the best reproducibility between replicates (CV = 14.8%) (Figure 3C).

**Figure 3.**
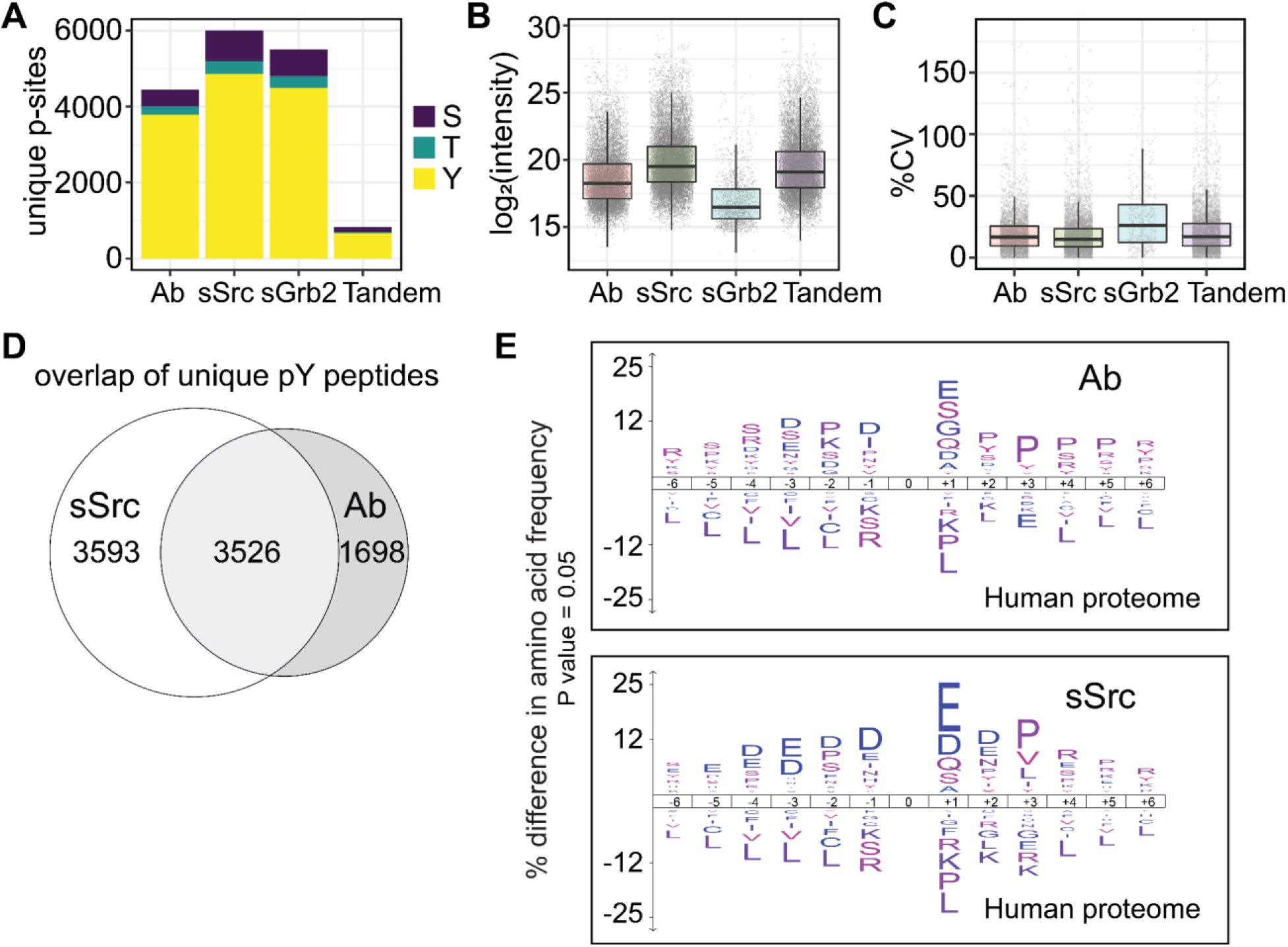
Different pY capture reagents display distinct sensitivity and reproducibility. **A.** Barplot depicting numbers of unique, quantifiable and confidently localized phosphosites across triplicate enrichments with each binder, colored by phosphorylated amino acid. **B**. Boxplot depicting log2 MS1 signal intensities of pY-peptides from triplicate enrichments with each binder. **C**. Boxplot depicting distributions of percent coefficient of variation of pY-peptide intensity between replicates, after median normalization. **D.** Overlap of unique pY-peptides captured by sSrc and antibody mixture (1:1 pTyr-100 and p-Tyr-1000). **E.** Difference in amino acid frequency surrounding the pY modification in peptides enriched by antibodies or sSrc relative to amino acid abundances in the human proteome. The motif plot was generated with IceLogo^35^ using a p-value cutoff of 0.05. The human proteome was sampled by IceLogo to generate background amino acid frequencies. Heights of amino acids represent their relative enrichment or depletion. Peptides identified in multiple replicates were multiply counted in a set to reflect redundancy in sequence preference.

Although sSrc is the best performing pY-capture reagent, other binders are able to recover pY sites not captured by sSrc. Specifically, of the 6201 unique pY sites enriched across all binders, sSrc did not capture 1345 or 22% of the pY sites (Supporting Figure 2C). Therefore, utilization of alternate pY capture reagents within R2-pY can expand the detectable set of pY sites. We also note that the poor performance of sGrb2 is in line with previous findings that report sGrb2 captured less pY-peptides than wild-type Src SH2^21^. Taken together, these results demonstrate that each sSrc, tandem, and anti-pY antibodies individually capture high numbers of pY sites and together offer complementary pY site selectivity.

### Characterizing sequence specificity of different pY capture reagents in R2-pY

Given the successful pY enrichment by both antibody and sSrc within the R2-pY platform, we sought to compare the pY-peptide pools captured with each reagent. Sequence specific biases of common anti-phosphotyrosine antibodies have been established^14,39^. Similarly, sSrc displays sequence preference reminiscent of the parent Src SH2 domain^14^. Upon inspection of intersections of pY-peptide sequences enriched by each binder, we observed moderate overlap (40% of unique pY-peptide sequences were detected by both sSrc and antibodies) (Figure 3D). This suggests that each binder displays sequence specific biases. To illustrate the amino acid preferences of sSrc relative to antibodies, we used the IceLogo tool^35^ to compare position-specific amino acid frequencies within the set of pY-peptides enriched by each binder relative to amino acid frequencies in the human proteome (Figure 3E). Peptide sequences were mapped back to the full protein and extended 6 amino acids on both sides of pY sites. Sites identified in multiple replicates were multiply counted in a set to reflect redundancy in sequence preference. Within the resulting logos, the heights of amino acids represent differences in amino acid frequency between the pool of enriched pY-peptides and the human proteome (Figure 3E). Indeed, moderate differences in sequence selectivity are observed between sSrc and antibody (Figure 3E). Specifically, sSrc preferred acidic aspartic and glutamic residues at positions −5 to +2 along with valine, leucine and isoleucine at position +3 in agreement with the parent Src SH2 domain binding preference^40–45^ (Figure 3E). Maximal difference in amino acid composition was 5% relative enrichment of glutamic acid at position +1 (Figure 3E). Broadly, the sequence preference of sSrc relative to antibody can be summarized as (E) - (D/E)-(E/D)-(D/P/S)-(D/E)-pY-(E/D/Q)-(D/E/N)-(P/V/L/I) (Figure 3E). This consensus sequence shares elements of the three previously published Src SH2 and sSrc SH2 recognition motifs^14,40–45^, but has increased preference for glutamic and aspartic acid at positions −1 and −2 along with proline at +3 (Figure 3E). These differences likely arise from differences of the biological sample as different sample matrices slightly alter consensus binding motifs^20,40,44^. Similarly, antibodies show sequence preference relative to sSrc. Increased preference for proline at positions +2 to +5 reflect known bias of the p-Tyr-100 antibody^39^. Further, antibodies show preference for serine, arginine and lysine at positions −4 to −2 as well as glycine at position +1 relative to sSrc. The consensus motif of antibody relative to the human proteome can be summarized as (S/R)-(D/S/E)-(P/K/S)-(D/I)-pY-(E/S/G/Q/D)-(P/Y)-(P)-(P/S)-(P)-(P/R) (Figure 3E). The consensus motif partially aligns with previously documented preferences of three anti-pY antibodies^39^. Partial alignment of sequence preference relative to previous studies may be due to differences in sample matrices and motif analysis workflows^14,39^.

The partial overlap of pY-peptide sequences and the slight differences in observed sequence specificity between sSrc and antibodies was not surprising^14,46,47^. In fact, this suggests that combining multiple pY capture reagents can expand the set of pY-peptides identified^20,48^. Conversely, use of specific sSH2 domains may focus enrichment to specific pathways^20,47^. Ultimately, the capture reagent(s) should remain consistent within an experiment for accurate comparisons.

### Assessing pY binding capacity of sSrc beads in R2-pY

We showed sSrc conjugated to NHS-activated magnetic beads is a highly efficient pY capture reagent within the R2-pY platform, providing a cost-effective alternative to antibodies. We next determined the pY-peptide binding capacity of sSrc beads in order to maximize pY-peptide capture while minimizing reagent use. We aimed to find a range of sSrc beads where the pY-peptide intensity and number of pY sites increased then approached a plateau. We tested four amounts of sSrc beads spanning a 16-fold difference while keeping peptide input amount constant at 1 mg in the R2-pY to R2-P2 workflow in technical triplicate. Phosphopeptide intensities increased between 2.1 nmol sSrc and 8.3 nmol sSrc then started to plateau between 8.3 to 16.6 nmol of sSrc (Figure 4A). The highest amount of beads used (16.6 nmol of sSrc, 1.6 mg beads) did not reach the maximum binding capacity of the KingFisher magnetic tips, as we did not observe bead deposition in the plates. This indicates that the intensity plateau observed at 16.6 nmol of sSrc represents depletion of pY-peptides from the sample instead of saturation of the magnetic tip capacity. Similarly, the number of unique pY sites increased between 2.1 nmol and 8.3 nmol of sSrc, then showed lesser increase between 8.3 nmol and 16.6 nmol of sSrc (Figure 4B). These results suggest that 8.3 nmol to 16.6 nmol of sSrc achieve near complete capture of the pY-peptides from 1 mg of peptide input. Ultimately, 8.3 nmol of sSrc provides reasonably comprehensive capture of the pY-peptides while minimizing reagent usage. Therefore we use 8.3 nmol of sSrc per 1 mg of HeLa peptides in subsequent experiments.

**Figure 4.**
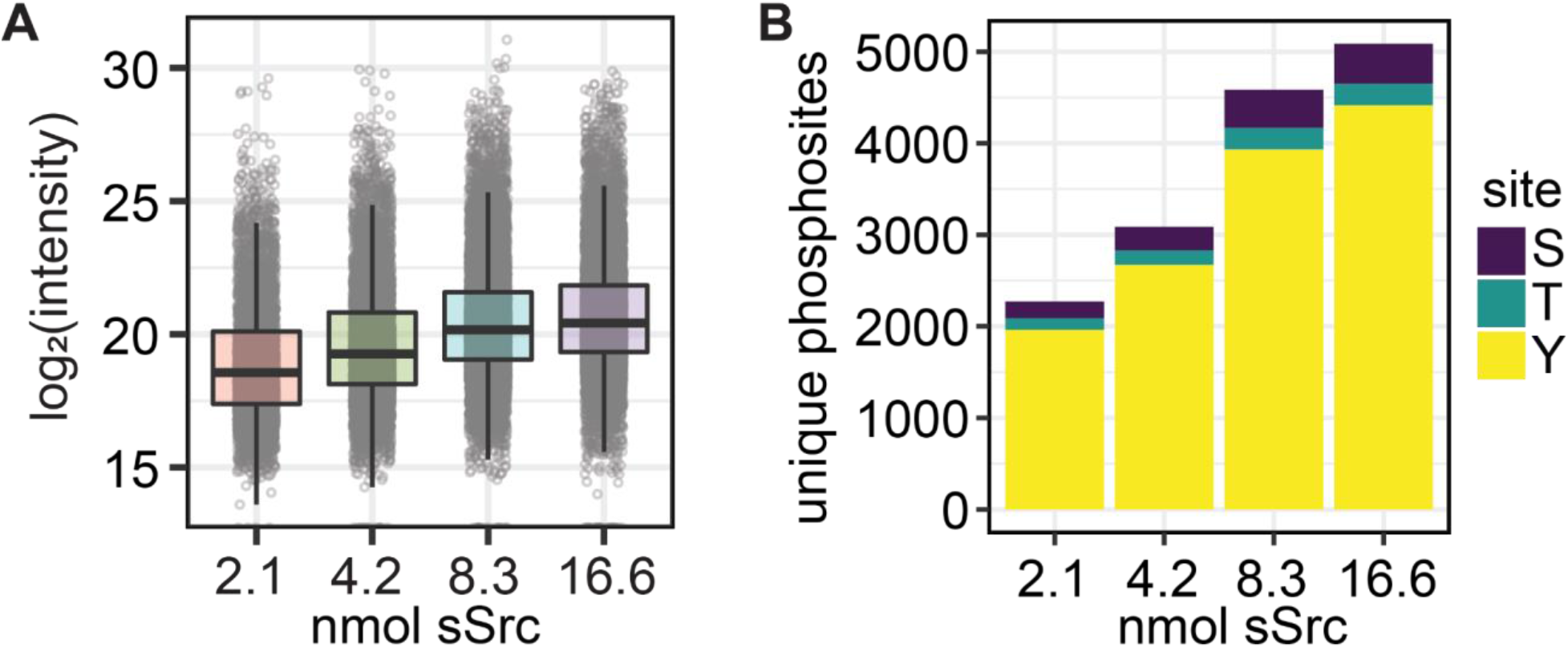
Comparison of sSrc amounts for complete pY-proteome enrichment. **A.** Boxplot depicting log2 MS1 precursor intensities of pY-peptides enriched with increasing amounts of sSrc and a fixed peptide amount (1mg). **B**. Barplot depicting numbers of unique phosphosites identified with increasing amounts of sSrc and a fixed peptide amount (1mg) in triplicate enrichments; colored by phosphorylated amino acid.

### Assessing linearity of pY capture by sSrc beads in R2-pY

A requirement for the quantitative use of a phosphotyrosine enrichment is linearity between quantified MS signal intensities and peptide input amounts. The ideal enrichment should allow detecting small amounts of phosphotyrosine peptides without saturating capture and detection in pY-rich samples. To characterize the quantitative scalability of R2-pY, we varied the peptide input amount a total of 16-fold using 0.25 mg, 1 mg and 4 mg of peptide input and conducted enrichments in technical triplicate. For this experiment we kept the bead to peptide ratio constant at 8.3 nmol sSrc per 1 mg of peptides, as we established earlier. The number of unique pY sites drastically increased between 0.25 mg and 1 mg peptide input from 2726 to 4739 pY sites, while increasing input amounts to 4 mg only provided marginal improvement (4937 unique pY sites) (Figure 5A). To determine the relationship between R2-pY-peptide input amount and resulting pY-peptide quantifications, we performed linear regressions on MS1 intensities for individual pY-peptides (Figure 5C and Supporting Figure 3). As expected, the majority of pY-peptide intensities increased linearly with input amounts (Figure 5B, 5C and Supporting Figure 3) in the tested range from 0.25 mg to 4 mg (Figure 5B and Supporting Figure 3) with a median slope of 1.14 (Figure 5C, left). Precision of linear regression models for each pY-peptide was assessed with Pearson’s R correlation coefficient. Median R^2^ was 0.96 and overall distribution of Pearson’s R^2^ for individual peptide intensities suggests good correlation of each peptide with its linear model (Figure 5C, right). Taken together, these results validate R2-pY for identifying thousands of pY sites and assessing their quantitative differences between samples.

**Figure 5.**
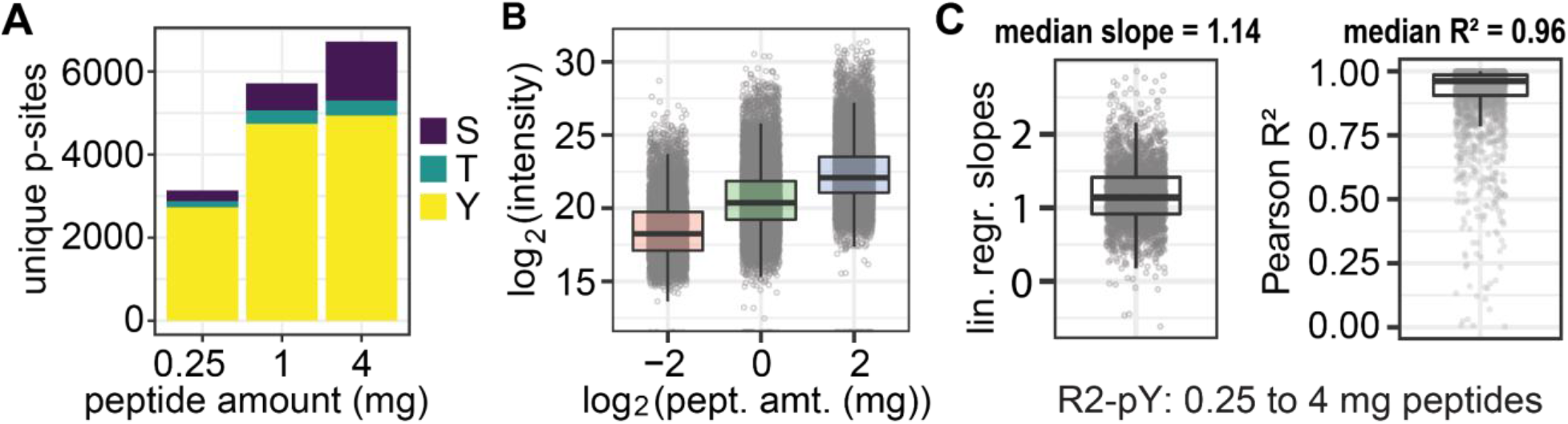
R2-pY is quantitatively scalable. **A.** Number of unique, quantifiable and confidently localized phosphosites for each peptide input amount, colored by phosphorylated amino acid. **B.** Boxplot depicting log_2_ intensity of peptides with confidently localized pY site(s) relative to log_2_ of peptide input amount. **C.** *Left:* Boxplot depicting distribution of linear regression slopes for each pY-peptide plotted in B (n = 2979). *Right:* Boxplot depicting distribution of Pearson R^2^ values demonstrating correlation of each pY-peptide intensity to its linear model across the three peptide input amounts (n = 2979).

### Integrating the R2-pY module in the R2-P2 automated workflow

With all the optimizations, R2-pY is a good standalone method for automated phosphotyrosine enrichment. However, R2-pY is also modular within our robotic pipeline; it can be easily integrated with previous R2-P2 protocols^28^ to streamline production of total proteome, global phosphoproteome and tyrosine-specific phosphoproteome peptide fractions (Figure 6A and Figure 1A-C). To demonstrate the performance of R2-pY within the integrated workflow, we processed lysates with R2-P1 instead of the in-solution digestion and solid-phase extraction desalting used for the optimization experiments described above. Subsequently pY-peptides were enriched using sSrc or p-Tyr-100:p-Tyr-1000 antibody mix magnetic beads in the R2-pY module and stringently cleaned by Fe^3+^-IMAC in the R2-P2 module prior to MS measurement by DDA. Enrichments were performed in technical duplicate from 1 mg of peptide input.

**Figure 6.**
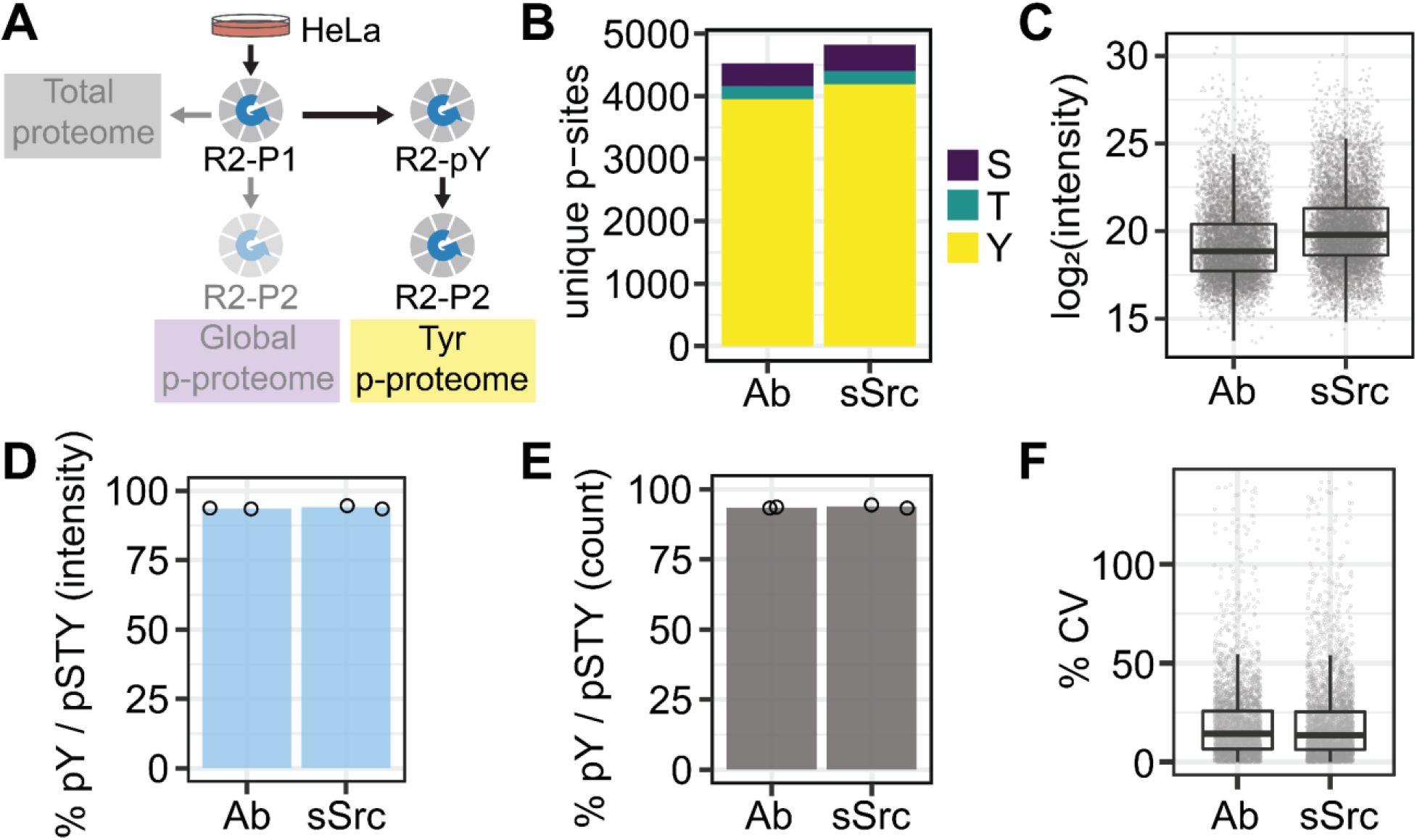
R2-pY functions as an integrated module within R2-P1 and R2-P2. **A.** Overview of workflow for validation of the fully automated pY enrichment module within the context of the full sample processing platform. **B.** Number of unique phosphosites identified by each binder after technical duplicate enrichments. **C.** Boxplot of MS1 signal intensities for unique pY-peptides enriched by either pTyr100:pTyr1000 antibody mix (Ab) or sSrc in R2-pY. Only peptides with at least one confidently localized pY site are plotted. **D.** pY-peptide enrichment efficiency as a ratio of precursor intensities. **E.** pY-peptide enrichment efficiency as a ratio of peptide count. **F.** Boxplot of percent coefficient of variation for pY-peptide intensities between replicates for both Ab and sSrc binders.

Results from the fully automated peptide preparation, pY-peptide enrichment, and cleanup were evaluated according to enrichment depth, distribution of phosphopeptide intensities, enrichment efficiency, and reproducibility between technical replicates. We show that similarly to the in-solution digestion and solid-phase extraction desalting, peptide preparation with R2-P1 also leads to efficient and reproducible pY-peptide enrichment with R2-pY to R2-P2. Under the tested conditions, R2-pY enrichment with sSrc captured pY-peptides slightly more efficiently than the pY antibody mixture, assessed by the number of unique pY sites identified (4187 vs. 3949 unique pY sites) (Figure 6A) and the higher MS1 precursor intensities (1.05 fold greater median log2(intensity)) (Figure 6B). Regardless of these differences, both binders achieved remarkably high phosphopeptide enrichment, with more than 99% of peptides identified and MS1 signal attributed to phosphopeptides and more than 93% of those peptides contain a confidently localized pY site (Supporting Figure 4A, B and Figure 6D, E). We also found good reproducibility between replicates assessed by the low coefficients of variation (CV) (below 15%) and high Pearson’s correlations (R^2^) (0.9 for sSrc and 0.9 for antibody) of pY-peptide intensity between technical duplicates (Figure 6F and Supporting Figure 4C). We note the amount of antibody was not optimized for this sample; we instead compared equal volumes of bead slurry. Therefore, optimization of antibody amounts may improve pY enrichment depth. This data demonstrates that R2-P1 works well for processing of lysates to peptides prior to R2-pY and that the fully automated workflow offers efficient and reproducible pY-peptide enrichment.

### R2-pY enables study of kinase phosphoregulation

The previous section demonstrates that R2-pY captures the tyrosine phosphoproteome efficiently and reproducibly in the context of our automated pipeline. Importantly, the datasets generated capture a number of proteins central to phosphorylation signaling. We highlight phosphosites observed on human kinases given the central importance of kinase activation in signal transduction and the association of kinase dysregulation with disease^49^. We report that after pY-peptide enrichment from 1 mg of peptide input in triplicate, antibody enrichment led to the identification of 128 unique pY sites across 64 human kinases and sSrc enrichment yielded 138 unique pY sites across 70 human kinases (Figure 7A). Notably, pY sites were detected on kinases across all kinase groups except casein kinases (CK1).

**Figure 7.**
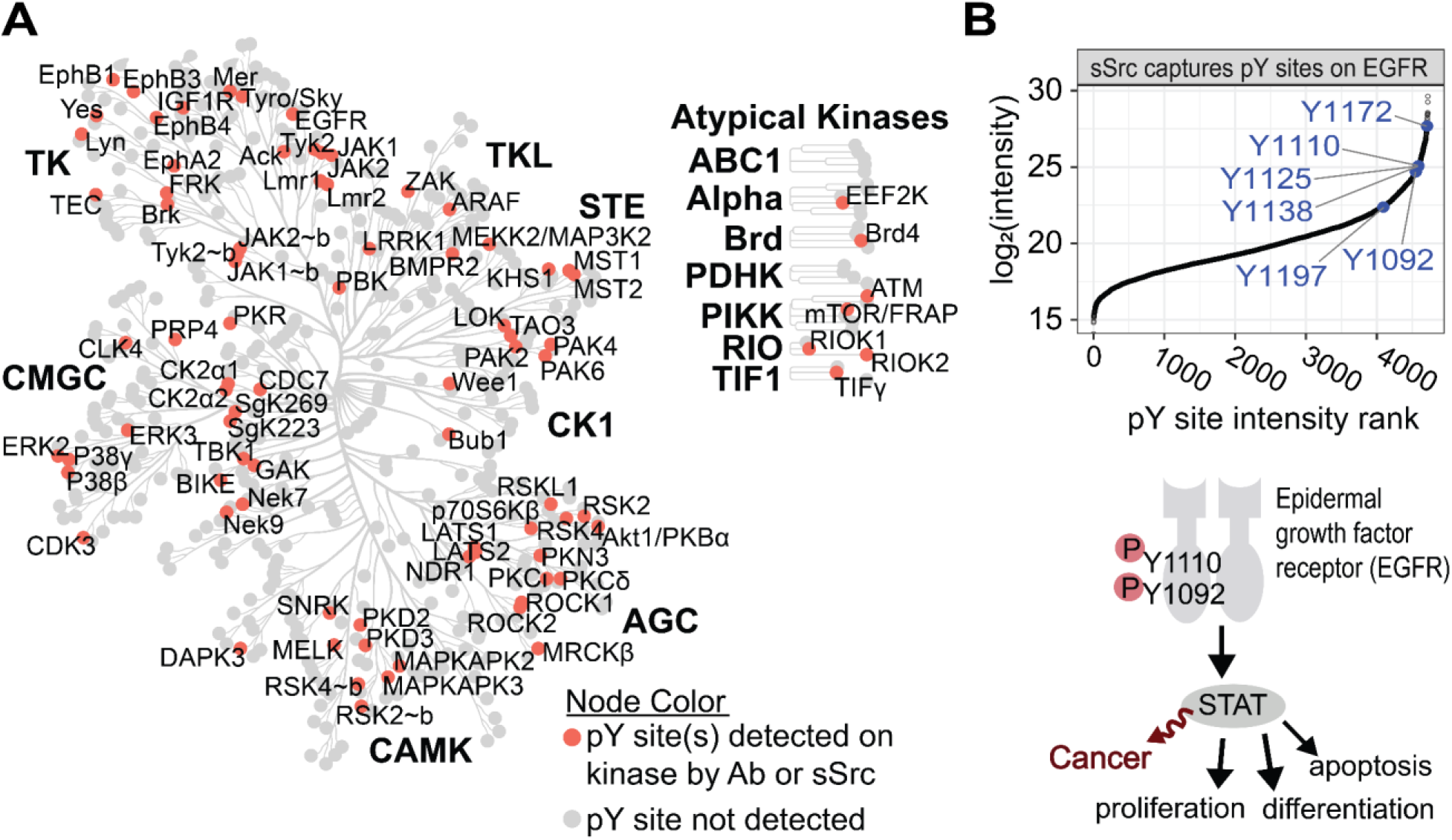
Examples of pY sites on human kinases detected from HeLa digest with R2-pY. **A.** Dendrogram of the human kinome that highlights kinases with confidently localized pY site(s) detected after R2-pY enrichment from HeLa cell proteins. **B**. *Top:* Intensity rank plot of six pY sites detected on epidermal growth factor receptor (EGFR) using R2-pY with sSrc beads. *Right:* Model highlighting two pY sites constitutively active in numerous cancers that are successfully captured by the R2-pY method.

Further, we highlight the sensitivity of R2-pY with sSrc beads to provide relative quantification of pY sites detected on epidermal growth factor receptor (EGFR) (Figure 7B). Six different pY sites were found on EGFR (Y1092, Y1110, Y1125, Y1138, Y1172, and Y1197) and measured at moderate to high intensity relative to other pY-peptides detected (Figure 7B). Some of these pY sites on EGFR promote STAT3 DNA binding activity and cell proliferation and have been found upregulated in several cancers^50^ (Figure 7B). R2-pY’s ability to efficiently capture biologically relevant pY sites facilitates their reproducible quantification by LC-MS/MS, enabling many applications in basic biology and translational research. Further, pY-profiling exemplified here can be achieved in 96 samples simultaneously opening the door to high-throughput and multidimensional signaling studies. In summary, we highlight capabilities of R2-pY to enrich a wide range of pY sites as illustrated by representation of pY sites on 7 of the 8 human kinase groups. We also note R2-pY enriches thousands of pY sites beyond those found on kinases thus enabling investigations of broader signaling networks.

## Conclusion

The R2-pY method presented here adds a new phosphotyrosine (pY) enrichment module to the R2-P2 workflow facilitating high dimensional pY signaling studies alongside global proteome and phosphoproteome profiling. This study optimized several key parameters of the new module to provide high sample purity, reproducibility, scalability, and versatility with regards to different pY capture reagents, including pY antibodies and SH2 superbinders. We validate the method with HeLa cell lysate and expect R2-pY to be amenable to other complex sample matrices. Further, while we evaluate R2-pY with data-dependent MS acquisition, samples are suitable for other MS-acquisition techniques. The robotic pY enrichment module will be attractive for any proteomic laboratory dealing with large sample batches, limited sample material and desire to reduce cost without sacrificing pY investigation depth. We have shown efficiency and reproducibility of R2-pY to enrich pY-peptides and anticipate the platform can be further expanded to study other modifications such as lysine acetylation, ubiquitylation and ADP ribosylation through the use of antibodies and binding domains^51–55^.

## Data availability

All mass spectrometry proteomics data have been deposited to the ProteomeXchange Consortium (http://www.proteomexchange.org/) via the PRIDE partner repository ^56^ with the dataset identifier PXD038788.

## Supporting Information

Supporting Methods 1. Standard operating procedure for R2-pY

Supporting Methods 2. Rationale for testing different conjugation chemistries

Supporting File 1. Superbinder sequences

Supporting File 2. Human protein sequence database

Supporting Figure 1. Example SDS PAGE for affinity purification of pY superbinders

Supporting Figure 2. Enrichment efficiency and overlap of different pY affinity reagents

Supporting Figure 3. R2-pY is quantitatively scalable

Supporting Figure 4. R2-P1 to R2-P2 to R2-pY displays high phosphopeptide enrichment efficiency.

Supporting Table 1A. All peptides from conjugation chemistry comparison.

Supporting Table 1B. Confidently localized phosphosites from conjugation chemistry comparison.

Supporting Table 1C. Ambiguous phosphosites from conjugation chemistry comparison.

Supporting Table 2A. All peptides from pY-peptide binding capacity tests of sSrc beads.

Supporting Table 2B. Confidently localized phosphosites from pY-peptide binding capacity tests of sSrc beads.

Supporting Table 2C. Ambiguous phosphosites from pY-peptide binding capacity tests of sSrc beads.

Supporting Table 3A. All peptides from peptide input scalability tests.

Supporting Table 3B. Confidently localized phosphosites from peptide input scalability tests.

Supporting Table 3C. Ambiguous phosphosites from peptide input scalability tests.

Supporting Table 4A. All peptides from pY affinity reagent comparison.

Supporting Table 4B. Confidently localized phosphosites from pY affinity reagent comparison.

Supporting Table 4C. Ambiguous phosphosites from pY affinity reagent comparison.

Supporting Table 5A. All peptides from tests of R2-P1 to R2-P2 to R2-pY.

Supporting Table 5B. Confidently localized phosphosites from tests of R2-P1 to R2-P2 to R2-pY.

Supporting Table 5C. Ambiguous phosphosites from tests of R2-P1 to R2-P2 to R2-pY.

## Acknowledgments

The authors would like to thank Ariadna Llovet and Julian Ramos for providing support with in-house proteomics software, and Sophie Moggridge, Matthew Berg, and former members of the Villen lab for providing valuable feedback on this work.

## Funding

This work was supported by NIH grants R35GM119536 and R01AG056359, and grant RGP0034/2018 from the Human Frontiers Science Program (to J.V.).

## Notes

http://www.ebi.ac.uk/pride

### Competing Interest Statement

The authors have declared no competing interest.

