## Supplemental Files for "Automated Enrichment of Phosphotyrosine Peptides for High-Throughput Proteomics"

#### Supporting Information

Supporting File 1. Superbinder sequences

Supporting Methods 1. Standard operating procedure for R2-pY

Supporting Methods 2. Rationale for testing different conjugation chemistries

Supporting Figure 1. Example SDS PAGE for affinity purification of pY superbinders

Supporting Figure 2. Enrichment efficiency and overlap of different pY affinity reagents

Supporting Figure 3. R2-pY is quantitatively scalable

Supporting Figure 4. R2-P1 to R2-P2 to R2-pY displays high phosphopeptide enrichment efficiency.

Supporting Table 1A. All peptides from conjugation chemistry comparison.

Supporting Table 1B. Confidently localized phosphosites from conjugation chemistry comparison.

Supporting Table 1C. Ambiguous phosphosites from conjugation chemistry comparison.

Supporting Table 2A. All peptides from pY-peptide binding capacity tests of sSrc beads.

Supporting Table 2B. Confidently localized phosphosites from pY-peptide binding capacity tests of sSrc beads.

Supporting Table 2C. Ambiguous phosphosites from pY-peptide binding capacity tests of sSrc beads.

Supporting Table 3A. All peptides from peptide input scalability tests.

Supporting Table 3B. Confidently localized phosphosites from peptide input scalability tests.

Supporting Table 3C. Ambiguous phosphosites from peptide input scalability tests.

Supporting Table 4A. All peptides from pY affinity reagent comparison.

Supporting Table 4B. Confidently localized phosphosites from pY affinity reagent comparison.

Supporting Table 4C. Ambiguous phosphosites from pY affinity reagent comparison.

Supporting Table 5A. All peptides from tests of R2-P1 to R2-P2 to R2-pY.

Supporting Table 5B. Confidently localized phosphosites from tests of R2-P1 to R2-P2 to R2-pY.

Supporting Table 5C. Ambiguous phosphosites from tests of R2-P1 to R2-P2 to R2-pY.

### Supporting File 1:

#### Superbinder sequences

**Construct name**: GRB2_SH2_A8V_S10A_K15L in pET28a(+)
**Full DNA sequence including plasmid**:
TAATACGACTCACTATAGGGGAATTGTGAGCGGATAACAATTCCCCTCTAGAAATAATTTTGTT
TAACTTTAAGAAGGAGATATACCATGGGCAGCAGCCATCATCATCATCATCACAGCAGCGGC
CTGGTGCCGCGCGGCAGCCATATGAAGCCTCATCCTTGGTTCTTTGGTAAGATCCCCCGTG
CGAAAGCTGAAGAGATGCTGTCAAAGCAACGTCACGACGGGGCCTTCCTTATCCGTGAGA
GTGAATCTGTTCCTGGTGACTTCGCTTTATCTGTAAAGTTCGGTAACGACGTGCAACATTTC
CTGGTGCTGCGCGATGGTGCGGGGAAGTACTTCTTATGGGTTGTTAAGTTCAATTCCTTGAA
TGAACTTGTAGACTATCATCGCAGCACCAGCGTGAGCCGCAACCAGCAAATCTTTCTCCGT
GATATTGAGCAGGTTCCGCAACAGCCTTGACTCGAGCACCACCACCACCACCACTGAGATC
CGGCTGCTAACAAAGCCCGAAAGGAAGCTGAGTTGGCTGCTGCCACCGCTGAGCAATAAC
TAGCATAACCCCTTGGGGCCTCTAAACGGGTCTTGAGGGGTTTTTTGCTGAAAGGAGGAAC
TATATCCGGATTGGCGAATGGGACGCGCCCTGTAGCGGCGCATTAAGCGCGGCGGGTGTG
GTGGTTACGCGCAGCGTGACCGCTACACTTGCCAGCGCCCTAGCGCCCGCTCCTTTCGCT
TTCTTCCCTTCCTTTCTCGCCACGTTCGCCGGCTTTCCCCGTCAAGCTCTAAATCGGGGGC
TCCCTTTAGGGTTCCGATTTAGTGCTTTACGGCACCTCGACCCCAAAAAACTTGATTAGGGT
GATGGTTCACGTAGTGGGCCATCGCCCTGATAGACGGTTTTTCGCCCTTTGACGTTGGAGT
CCACGTTCTTTAATAGTGGACTCTTGTTCCAAACTGGAACAACACTCAACCCTATCTCGGTC
TATTCTTTTGATTTATAAGGGATTTTGCCGATTTCGGCCTATTGGTTAAAAAATGAGCTGATTTA
ACAAAAATTTAACGCGAATTTTAACAAAATATTAACGCTTACAATTTAGGTGGCACTTTTCGGG
GAAATGTGCGCGGAACCCCTATTTGTTTATTTTTCTAAATACATTCAAATATGTATCCGCTCAT
GAATTAATTCTTAGAAAAACTCATCGAGCATCAAATGAAACTGCAATTTATTCATATCAGGATTA
TCAATACCATATTTTTGAAAAAGCCGTTTCTGTAATGAAGGAGAAAACTCACCGAGGCAGTTC
CATAGGATGGCAAGATCCTGGTATCGGTCTGCGATTCCGACTCGTCCAACATCAATACAACC
TATTAATTTCCCCTCGTCAAAAATAAGGTTATCAAGTGAGAAATCACCATGAGTGACGACTGA
ATCCGGTGAGAATGGCAAAAGTTTATGCATTTCTTTCCAGACTTGTTCAACAGGCCAGCCAT
TACGCTCGTCATCAAAATCACTCGCATCAACCAAACCGTTATTCATTCGTGATTGCGCCTGA
GCGAGACGAAATACGCGATCGCTGTTAAAAGGACAATTACAAACAGGAATCGAATGCAACC
GGCGCAGGAACACTGCCAGCGCATCAACAATATTTTCACCTGAATCAGGATATTCTTCTAATA
CCTGGAATGCTGTTTTCCCGGGGATCGCAGTGGTGAGTAACCATGCATCATCAGGAGTACG
GATAAAATGCTTGATGGTCGGAAGAGGCATAAATTCCGTCAGCCAGTTTAGTCTGACCATCT
CATCTGTAACATCATTGGCAACGCTACCTTTGCCATGTTTCAGAAACAACTCTGGCGCATCG
GGCTTCCCATACAATCGATAGATTGTCGCACCTGATTGCCCGACATTATCGCGAGCCCATTTA
TACCCATATAAATCAGCATCCATGTTGGAATTTAATCGCGGCCTAGAGCAAGACGTTTCCCGT
TGAATATGGCTCATAACACCCCTTGTATTACTGTTTATGTAAGCAGACAGTTTTATTGTTCATG
ACCAAAATCCCTTAACGTGAGTTTTCGTTCCACTGAGCGTCAGACCCCGTAGAAAAGATCAA
AGGATCTTCTTGAGATCCTTTTTTTCTGCGCGTAATCTGCTGCTTGCAAACAAAAAAACCAC
CGCTACCAGCGGTGGTTTGTTTGCCGGATCAAGAGCTACCAACTCTTTTTCCGAAGGTAAC

TGGCTTCAGCAGAGCGCAGATACCAAATACTGTCCTTCTAGTGTAGCCGTAGTTAGGCCACC
ACTTCAAGAACTCTGTAGCACCGCCTACATACCTCGCTCTGCTAATCCTGTTACCAGTGGCT
GCTGCCAGTGGCGATAAGTCGTGTCTTACCGGGTTGGACTCAAGACGATAGTTACCGGATA
AGGCGCAGCGGTCGGGCTGAACGGGGGGTTCGTGCACACAGCCCAGCTTGGAGCGAAC
GACCTACACCGAACTGAGATACCTACAGCGTGAGCTATGAGAAAGCGCCACGCTTCCCGAA
GGGAGAAAGGCGGACAGGTATCCGGTAAGCGGCAGGGTCGGAACAGGAGAGCGCACGAG
GGAGCTTCCAGGGGGAAACGCCTGGTATCTTTATAGTCCTGTCGGGTTTCGCCACCTCTGA
CTTGAGCGTCGATTTTTGTGATGCTCGTCAGGGGGGCGGAGCCTATGGAAAAACGCCAGC
AACGCGGCCTTTTTACGGTTCCTGGCCTTTTGCTGGCCTTTTGCTCACATGTTCTTTCCTGC
GTTATCCCCTGATTCTGTGGATAACCGTATTACCGCCTTTGAGTGAGCTGATACCGCTCGCC
GCAGCCGAACGACCGAGCGCAGCGAGTCAGTGAGCGAGGAAGCGGAAGAGCGCCTGATG
CGGTATTTTCTCCTTACGCATCTGTGCGGTATTTCACACCGCAATGGTGCACTCTCAGTACA
ATCTGCTCTGATGCCGCATAGTTAAGCCAGTATACACTCCGCTATCGCTACGTGACTGGGTC
ATGGCTGCGCCCCGACACCCGCCAACACCCGCTGACGCGCCCTGACGGGCTTGTCTGCT
CCCGGCATCCGCTTACAGACAAGCTGTGACCGTCTCCGGGAGCTGCATGTGTCAGAGGTT
TTCACCGTCATCACCGAAACGCGCGAGGCAGCTGCGGTAAAGCTCATCAGCGTGGTCGTG
AAGCGATTCACAGATGTCTGCCTGTTCATCCGCGTCCAGCTCGTTGAGTTTCTCCAGAAGC
GTTAATGTCTGGCTTCTGATAAAGCGGGCCATGTTAAGGGCGGTTTTTTCCTGTTTGGTCAC
TGATGCCTCCGTGTAAGGGGGATTTCTGTTCATGGGGGTAATGATACCGATGAAACGAGAG
AGGATGCTCACGATACGGGTTACTGATGATGAACATGCCCGGTTACTGGAACGTTGTGAGG
GTAAACAACTGGCGGTATGGATGCGGCGGGACCAGAGAAAAATCACTCAGGGTCAATGCCA
GCGCTTCGTTAATACAGATGTAGGTGTTCCACAGGGTAGCCAGCAGCATCCTGCGATGCAG
ATCCGGAACATAATGGTGCAGGGCGCTGACTTCCGCGTTTCCAGACTTTACGAAACACGGA
AACCGAAGACCATTCATGTTGTTGCTCAGGTCGCAGACGTTTTGCAGCAGCAGTCGCTTCA
CGTTCGCTCGCGTATCGGTGATTCATTCTGCTAACCAGTAAGGCAACCCCGCCAGCCTAGC
CGGGTCCTCAACGACAGGAGCACGATCATGCGCACCCGTGGGGCCGCCATGCCGGCGAT
AATGGCCTGCTTCTCGCCGAAACGTTTGGTGGCGGGACCAGTGACGAAGGCTTGAGCGAG
GGCGTGCAAGATTCCGAATACCGCAAGCGACAGGCCGATCATCGTCGCGCTCCAGCGAAA
GCGGTCCTCGCCGAAAATGACCCAGAGCGCTGCCGGCACCTGTCCTACGAGTTGCATGAT
AAAGAAGACAGTCATAAGTGCGGCGACGATAGTCATGCCCCGCGCCCACCGGAAGGAGCT
GACTGGGTTGAAGGCTCTCAAGGGCATCGGTCGAGATCCCGGTGCCTAATGAGTGAGCTA
ACTTACATTAATTGCGTTGCGCTCACTGCCCGCTTTCCAGTCGGGAAACCTGTCGTGCCAG
CTGCATTAATGAATCGGCCAACGCGCGGGGAGAGGCGGTTTGCGTATTGGGCGCCAGGGT
GGTTTTTCTTTTCACCAGTGAGACGGGCAACAGCTGATTGCCCTTCACCGCCTGGCCCTGA
GAGAGTTGCAGCAAGCGGTCCACGCTGGTTTGCCCCAGCAGGCGAAAATCCTGTTTGATG
GTGGTTAACGGCGGGATATAACATGAGCTGTCTTCGGTATCGTCGTATCCCACTACCGAGAT
ATCCGCACCAACGCGCAGCCCGGACTCGGTAATGGCGCGCATTGCGCCCAGCGCCATCTG
ATCGTTGGCAACCAGCATCGCAGTGGGAACGATGCCCTCATTCAGCATTTGCATGGTTTGTT
GAAAACCGGACATGGCACTCCAGTCGCCTTCCCGTTCCGCTATCGGCTGAATTTGATTGCG
AGTGAGATATTTATGCCAGCCAGCCAGACGCAGACGCGCCGAGACAGAACTTAATGGGCCC
GCTAACAGCGCGATTTGCTGGTGACCCAATGCGACCAGATGCTCCACGCCCAGTCGCGTA
CCGTCTTCATGGGAGAAAATAATACTGTTGATGGGTGTCTGGTCAGAGACATCAAGAAATAA
CGCCGGAACATTAGTGCAGGCAGCTTCCACAGCAATGGCATCCTGGTCATCCAGCGGATAG
TTAATGATCAGCCCACTGACGCGTTGCGCGAGAAGATTGTGCACCGCCGCTTTACAGGCTT

CGACGCCGCTTCGTTCTACCATCGACACCACCACGCTGGCACCCAGTTGATCGGCGCGAG
ATTTAATCGCCGCGACAATTTGCGACGGCGCGTGCAGGGCCAGACTGGAGGTGGCAACGC
CAATCAGCAACGACTGTTTGCCCGCCAGTTGTTGTGCCACGCGGTTGGGAATGTAATTCAG
CTCCGCCATCGCCGCTTCCACTTTTTCCCGCGTTTTCGCAGAAACGTGGCTGGCCTGGTTC
ACCACGCGGGAAACGGTCTGATAAGAGACACCGGCATACTCTGCGACATCGTATAACGTTA
CTGGTTTCACATTCACCACCCTGAATTGACTCTCTTCCGGGCGCTATCATGCCATACCGCGA
AAGGTTTTGCGCCATTCGATGGTGTCCGGGATCTCGACGCTCTCCCTTATGCGACTCCTGC
ATTAGGAAGCAGCCCAGTAGTAGGTTGAGGCCGTTGAGCACCGCCGCCGCAAGGAATGGT
GCATGCAAGGAGATGGCGCCCAACAGTCCCCCGGCCACGGGGCCTGCCACCATACCCAC
GCCGAAACAAGCGCTCATGAGCCCGAAGTGGCGAGCCCGATCTTCCCCATCGGTGATGTC
GGCGATATAGGCGCCAGCAACCGCACCTGTGGCGCCGGTGATGCCGGCCACGATGCGTC
CGGCGTAGAGGATCGAGATCTCGATCCCGCGAAAT

**Protein sequence (CDS):**
MGSSHHHHHHSSGLVPRGSHMKPHPWFFGKIPRAKAEEMLSKQRHDGAFLIRESESVPGDFA
LSVKFGNDVQHFLVLRDGAGKYFLWVVKFNSLNELVDYHRSTSVSRNQQIFLRDIEQVPQQP*

**Construct name**: Src_SH2_T8V_C10A_K15L in pET28a(+)
**Full DNA sequence including plasmid**:
TAATACGACTCACTATAGGGGAATTGTGAGCGGATAACAATTCCCCTCTAGAAATAATTTTGTT
TAACTTTAAGAAGGAGATATACCATGGGCAGCAGCCATCATCATCATCATCACAGCAGCGGC
CTGGTGCCGCGCGGCAGCCATATGGACTCTATCCAAGCCGAGGAATGGTACTTTGGGAAAA
TCACGCGTCGTGAGTCCGAGCGTCTTCTCCTGAATGCGGAAAATCCGCGCGGCACCTTCTT
AGTTCGTGAGAGCGAGACAGTGAAGGGAGCCTACGCATTAAGCGTGAGCGACTTCGATAAT
GCCAAGGGTCTTAACGTAAAGCATTATTTGATTCGCAAACTCGACTCGGGCGGCTTCTATATT
ACTTCGCGTACTCAGTTCAACTCTCTCCAACAGTTAGTAGCATATTATTCTAAGCACGCGGAT
GGTCTGTGTCACCGCCTGACTACGGTTTGTCCCACTAGTAAGTGACTCGAGCACCACCACC
ACCACCACTGAGATCCGGCTGCTAACAAAGCCCGAAAGGAAGCTGAGTTGGCTGCTGCCA
CCGCTGAGCAATAACTAGCATAACCCCTTGGGGCCTCTAAACGGGTCTTGAGGGGTTTTTT
GCTGAAAGGAGGAACTATATCCGGATTGGCGAATGGGACGCGCCCTGTAGCGGCGCATTAA
GCGCGGCGGGTGTGGTGGTTACGCGCAGCGTGACCGCTACACTTGCCAGCGCCCTAGCG
CCCGCTCCTTTCGCTTTCTTCCCTTCCTTTCTCGCCACGTTCGCCGGCTTTCCCCGTCAAG
CTCTAAATCGGGGGCTCCCTTTAGGGTTCCGATTTAGTGCTTTACGGCACCTCGACCCCAAA
AAACTTGATTAGGGTGATGGTTCACGTAGTGGGCCATCGCCCTGATAGACGGTTTTTCGCC
CTTTGACGTTGGAGTCCACGTTCTTTAATAGTGGACTCTTGTTCCAAACTGGAACAACACTC
AACCCTATCTCGGTCTATTCTTTTGATTTATAAGGGATTTTGCCGATTTCGGCCTATTGGTTAA
AAAATGAGCTGATTTAACAAAAATTTAACGCGAATTTTAACAAAATATTAACGCTTACAATTTAG
GTGGCACTTTTCGGGGAAATGTGCGCGGAACCCCTATTTGTTTATTTTTCTAAATACATTCAA
ATATGTATCCGCTCATGAATTAATTCTTAGAAAAACTCATCGAGCATCAAATGAAACTGCAATT
TATTCATATCAGGATTATCAATACCATATTTTTGAAAAAGCCGTTTCTGTAATGAAGGAGAAAA
CTCACCGAGGCAGTTCCATAGGATGGCAAGATCCTGGTATCGGTCTGCGATTCCGACTCGT
CCAACATCAATACAACCTATTAATTTCCCCTCGTCAAAAATAAGGTTATCAAGTGAGAAATCAC
CATGAGTGACGACTGAATCCGGTGAGAATGGCAAAAGTTTATGCATTTCTTTCCAGACTTGT
TCAACAGGCCAGCCATTACGCTCGTCATCAAAATCACTCGCATCAACCAAACCGTTATTCATT
CGTGATTGCGCCTGAGCGAGACGAAATACGCGATCGCTGTTAAAAGGACAATTACAAACAG
GAATCGAATGCAACCGGCGCAGGAACACTGCCAGCGCATCAACAATATTTTCACCTGAATCA
GGATATTCTTCTAATACCTGGAATGCTGTTTTCCCGGGGATCGCAGTGGTGAGTAACCATGC
ATCATCAGGAGTACGGATAAAATGCTTGATGGTCGGAAGAGGCATAAATTCCGTCAGCCAGT
TTAGTCTGACCATCTCATCTGTAACATCATTGGCAACGCTACCTTTGCCATGTTTCAGAAACA
ACTCTGGCGCATCGGGCTTCCCATACAATCGATAGATTGTCGCACCTGATTGCCCGACATTA
TCGCGAGCCCATTTATACCCATATAAATCAGCATCCATGTTGGAATTTAATCGCGGCCTAGAG
CAAGACGTTTCCCGTTGAATATGGCTCATAACACCCCTTGTATTACTGTTTATGTAAGCAGAC
AGTTTTATTGTTCATGACCAAAATCCCTTAACGTGAGTTTTCGTTCCACTGAGCGTCAGACCC
CGTAGAAAAGATCAAAGGATCTTCTTGAGATCCTTTTTTTCTGCGCGTAATCTGCTGCTTGCA
AACAAAAAAACCACCGCTACCAGCGGTGGTTTGTTTGCCGGATCAAGAGCTACCAACTCTT
TTTCCGAAGGTAACTGGCTTCAGCAGAGCGCAGATACCAAATACTGTCCTTCTAGTGTAGCC
GTAGTTAGGCCACCACTTCAAGAACTCTGTAGCACCGCCTACATACCTCGCTCTGCTAATCC
TGTTACCAGTGGCTGCTGCCAGTGGCGATAAGTCGTGTCTTACCGGGTTGGACTCAAGACG
ATAGTTACCGGATAAGGCGCAGCGGTCGGGCTGAACGGGGGGTTCGTGCACACAGCCCAG
CTTGGAGCGAACGACCTACACCGAACTGAGATACCTACAGCGTGAGCTATGAGAAAGCGCC
ACGCTTCCCGAAGGGAGAAAGGCGGACAGGTATCCGGTAAGCGGCAGGGTCGGAACAGG

AGAGCGCACGAGGGAGCTTCCAGGGGGAAACGCCTGGTATCTTTATAGTCCTGTCGGGTTT
CGCCACCTCTGACTTGAGCGTCGATTTTTGTGATGCTCGTCAGGGGGGCGGAGCCTATGG
AAAAACGCCAGCAACGCGGCCTTTTTACGGTTCCTGGCCTTTTGCTGGCCTTTTGCTCACA
TGTTCTTTCCTGCGTTATCCCCTGATTCTGTGGATAACCGTATTACCGCCTTTGAGTGAGCTG
ATACCGCTCGCCGCAGCCGAACGACCGAGCGCAGCGAGTCAGTGAGCGAGGAAGCGGAA
GAGCGCCTGATGCGGTATTTTCTCCTTACGCATCTGTGCGGTATTTCACACCGCAATGGTGC
ACTCTCAGTACAATCTGCTCTGATGCCGCATAGTTAAGCCAGTATACACTCCGCTATCGCTAC
GTGACTGGGTCATGGCTGCGCCCCGACACCCGCCAACACCCGCTGACGCGCCCTGACGG
GCTTGTCTGCTCCCGGCATCCGCTTACAGACAAGCTGTGACCGTCTCCGGGAGCTGCATG
TGTCAGAGGTTTTCACCGTCATCACCGAAACGCGCGAGGCAGCTGCGGTAAAGCTCATCA
GCGTGGTCGTGAAGCGATTCACAGATGTCTGCCTGTTCATCCGCGTCCAGCTCGTTGAGTT
TCTCCAGAAGCGTTAATGTCTGGCTTCTGATAAAGCGGGCCATGTTAAGGGCGGTTTTTTCC
TGTTTGGTCACTGATGCCTCCGTGTAAGGGGGATTTCTGTTCATGGGGGTAATGATACCGAT
GAAACGAGAGAGGATGCTCACGATACGGGTTACTGATGATGAACATGCCCGGTTACTGGAA
CGTTGTGAGGGTAAACAACTGGCGGTATGGATGCGGCGGGACCAGAGAAAAATCACTCAG
GGTCAATGCCAGCGCTTCGTTAATACAGATGTAGGTGTTCCACAGGGTAGCCAGCAGCATC
CTGCGATGCAGATCCGGAACATAATGGTGCAGGGCGCTGACTTCCGCGTTTCCAGACTTTA
CGAAACACGGAAACCGAAGACCATTCATGTTGTTGCTCAGGTCGCAGACGTTTTGCAGCAG
CAGTCGCTTCACGTTCGCTCGCGTATCGGTGATTCATTCTGCTAACCAGTAAGGCAACCCC
GCCAGCCTAGCCGGGTCCTCAACGACAGGAGCACGATCATGCGCACCCGTGGGGCCGCC
ATGCCGGCGATAATGGCCTGCTTCTCGCCGAAACGTTTGGTGGCGGGACCAGTGACGAAG
GCTTGAGCGAGGGCGTGCAAGATTCCGAATACCGCAAGCGACAGGCCGATCATCGTCGCG
CTCCAGCGAAAGCGGTCCTCGCCGAAAATGACCCAGAGCGCTGCCGGCACCTGTCCTACG
AGTTGCATGATAAAGAAGACAGTCATAAGTGCGGCGACGATAGTCATGCCCCGCGCCCACC
GGAAGGAGCTGACTGGGTTGAAGGCTCTCAAGGGCATCGGTCGAGATCCCGGTGCCTAAT
GAGTGAGCTAACTTACATTAATTGCGTTGCGCTCACTGCCCGCTTTCCAGTCGGGAAACCT
GTCGTGCCAGCTGCATTAATGAATCGGCCAACGCGCGGGGAGAGGCGGTTTGCGTATTGG
GCGCCAGGGTGGTTTTTCTTTTCACCAGTGAGACGGGCAACAGCTGATTGCCCTTCACCG
CCTGGCCCTGAGAGAGTTGCAGCAAGCGGTCCACGCTGGTTTGCCCCAGCAGGCGAAAAT
CCTGTTTGATGGTGGTTAACGGCGGGATATAACATGAGCTGTCTTCGGTATCGTCGTATCCC
ACTACCGAGATATCCGCACCAACGCGCAGCCCGGACTCGGTAATGGCGCGCATTGCGCCC
AGCGCCATCTGATCGTTGGCAACCAGCATCGCAGTGGGAACGATGCCCTCATTCAGCATTT
GCATGGTTTGTTGAAAACCGGACATGGCACTCCAGTCGCCTTCCCGTTCCGCTATCGGCTG
AATTTGATTGCGAGTGAGATATTTATGCCAGCCAGCCAGACGCAGACGCGCCGAGACAGAA
CTTAATGGGCCCGCTAACAGCGCGATTTGCTGGTGACCCAATGCGACCAGATGCTCCACGC
CCAGTCGCGTACCGTCTTCATGGGAGAAAATAATACTGTTGATGGGTGTCTGGTCAGAGAC
ATCAAGAAATAACGCCGGAACATTAGTGCAGGCAGCTTCCACAGCAATGGCATCCTGGTCAT
CCAGCGGATAGTTAATGATCAGCCCACTGACGCGTTGCGCGAGAAGATTGTGCACCGCCG
CTTTACAGGCTTCGACGCCGCTTCGTTCTACCATCGACACCACCACGCTGGCACCCAGTTG
ATCGGCGCGAGATTTAATCGCCGCGACAATTTGCGACGGCGCGTGCAGGGCCAGACTGGA
GGTGGCAACGCCAATCAGCAACGACTGTTTGCCCGCCAGTTGTTGTGCCACGCGGTTGGG
AATGTAATTCAGCTCCGCCATCGCCGCTTCCACTTTTTCCCGCGTTTTCGCAGAAACGTGGC
TGGCCTGGTTCACCACGCGGGAAACGGTCTGATAAGAGACACCGGCATACTCTGCGACATC
GTATAACGTTACTGGTTTCACATTCACCACCCTGAATTGACTCTCTTCCGGGCGCTATCATGC

CATACCGCGAAAGGTTTTGCGCCATTCGATGGTGTCCGGGATCTCGACGCTCTCCCTTATG
CGACTCCTGCATTAGGAAGCAGCCCAGTAGTAGGTTGAGGCCGTTGAGCACCGCCGCCGC
AAGGAATGGTGCATGCAAGGAGATGGCGCCCAACAGTCCCCCGGCCACGGGGCCTGCCA
CCATACCCACGCCGAAACAAGCGCTCATGAGCCCGAAGTGGCGAGCCCGATCTTCCCCAT
CGGTGATGTCGGCGATATAGGCGCCAGCAACCGCACCTGTGGCGCCGGTGATGCCGGCCA
CGATGCGTCCGGCGTAGAGGATCGAGATCTCGATCCCGCGAAAT

**Protein sequence (CDS):**
MGSSHHHHHHSSGLVPRGSHMDSIQAEEWYFGKITRRESERLLLNAENPRGTFLVRESETVKG
AYALSVSDFDNAKGLNVKHYLIRKLDSGGFYITSRTQFNSLQQLVAYYSKHADGLCHRLTTVCPT
SK*

**Construct name**: tandemSH2_Src_Grb2 in pET28a(+)
**Full DNA sequence including plasmid**:
TAATACGACTCACTATAGGGGAATTGTGAGCGGATAACAATTCCCCTCTAGAAATAATTTTGTT
TAACTTTAAGAAGGAGATATACCATGGGCAGCAGCCATCATCATCATCATCACAGCAGCGGC
CTGGTGCCGCGCGGCAGCCATATGGACTCAATCCAGGCGGAAGAATGGTATTTCGGCAAAA
TTACCCGTCGCGAGTCTGAGCGCCTGCTCTTGAACGCTGAGAATCCGCGCGGAACTTTTCT
GGTCCGCGAGAGTGAGACAGTCAAAGGTGCCTATGCTCTGAGCGTCAGCGATTTTGATAAT
GCGAAGGGCTTAAACGTGAAACACTACTTAATCCGTAAGTTAGACTCGGGCGGCTTCTACAT
TACCTCGCGCACCCAGTTCAATAGCTTACAACAACTGGTAGCCTACTACTCCAAGCACGCAG
ATGGATTATGCCACCGCCTGACGACCGTATGTCCCACGTCTAAAATGAAGCCCCACCCCTG
GTTCTTCGGCAAAATTCCGCGTGCCAAGGCAGAAGAAATGCTGTCCAAGCAACGCCATGAC
GGCGCCTTCTTAATCCGCGAGAGCGAGAGCGTTCCTGGTGACTTCGCGTTATCGGTAAAAT
TTGGAAACGATGTCCAACATTTTCTGGTATTACGTGACGGTGCGGGCAAGTACTTCTTATGG
GTGGTAAAATTTAATTCTCTCAATGAGCTCGTAGACTACCACCGTAGCACTAGCGTCAGCCG
CAATCAACAGATTTTCCTTCGCGACATCGAGCAGGTACCGCAGCAACCATAACTCGAGCAC
CACCACCACCACCACTGAGATCCGGCTGCTAACAAAGCCCGAAAGGAAGCTGAGTTGGCT
GCTGCCACCGCTGAGCAATAACTAGCATAACCCCTTGGGGCCTCTAAACGGGTCTTGAGGG
GTTTTTTGCTGAAAGGAGGAACTATATCCGGATTGGCGAATGGGACGCGCCCTGTAGCGGC
GCATTAAGCGCGGCGGGTGTGGTGGTTACGCGCAGCGTGACCGCTACACTTGCCAGCGCC
CTAGCGCCCGCTCCTTTCGCTTTCTTCCCTTCCTTTCTCGCCACGTTCGCCGGCTTTCCCC
GTCAAGCTCTAAATCGGGGGCTCCCTTTAGGGTTCCGATTTAGTGCTTTACGGCACCTCGA
CCCCAAAAAACTTGATTAGGGTGATGGTTCACGTAGTGGGCCATCGCCCTGATAGACGGTT
TTTCGCCCTTTGACGTTGGAGTCCACGTTCTTTAATAGTGGACTCTTGTTCCAAACTGGAAC
AACACTCAACCCTATCTCGGTCTATTCTTTTGATTTATAAGGGATTTTGCCGATTTCGGCCTAT
TGGTTAAAAAATGAGCTGATTTAACAAAAATTTAACGCGAATTTTAACAAAATATTAACGCTTA
CAATTTAGGTGGCACTTTTCGGGGAAATGTGCGCGGAACCCCTATTTGTTTATTTTTCTAAAT
ACATTCAAATATGTATCCGCTCATGAATTAATTCTTAGAAAAACTCATCGAGCATCAAATGAAA
CTGCAATTTATTCATATCAGGATTATCAATACCATATTTTTGAAAAAGCCGTTTCTGTAATGAAG
GAGAAAACTCACCGAGGCAGTTCCATAGGATGGCAAGATCCTGGTATCGGTCTGCGATTCC
GACTCGTCCAACATCAATACAACCTATTAATTTCCCCTCGTCAAAAATAAGGTTATCAAGTGA
GAAATCACCATGAGTGACGACTGAATCCGGTGAGAATGGCAAAAGTTTATGCATTTCTTTCC
AGACTTGTTCAACAGGCCAGCCATTACGCTCGTCATCAAAATCACTCGCATCAACCAAACCG
TTATTCATTCGTGATTGCGCCTGAGCGAGACGAAATACGCGATCGCTGTTAAAAGGACAATT
ACAAACAGGAATCGAATGCAACCGGCGCAGGAACACTGCCAGCGCATCAACAATATTTTCA
CCTGAATCAGGATATTCTTCTAATACCTGGAATGCTGTTTTCCCGGGGATCGCAGTGGTGAG
TAACCATGCATCATCAGGAGTACGGATAAAATGCTTGATGGTCGGAAGAGGCATAAATTCCG
TCAGCCAGTTTAGTCTGACCATCTCATCTGTAACATCATTGGCAACGCTACCTTTGCCATGTT
TCAGAAACAACTCTGGCGCATCGGGCTTCCCATACAATCGATAGATTGTCGCACCTGATTGC
CCGACATTATCGCGAGCCCATTTATACCCATATAAATCAGCATCCATGTTGGAATTTAATCGCG
GCCTAGAGCAAGACGTTTCCCGTTGAATATGGCTCATAACACCCCTTGTATTACTGTTTATGT
AAGCAGACAGTTTTATTGTTCATGACCAAAATCCCTTAACGTGAGTTTTCGTTCCACTGAGC
GTCAGACCCCGTAGAAAAGATCAAAGGATCTTCTTGAGATCCTTTTTTTCTGCGCGTAATCT
GCTGCTTGCAAACAAAAAAACCACCGCTACCAGCGGTGGTTTGTTTGCCGGATCAAGAGCT
ACCAACTCTTTTTCCGAAGGTAACTGGCTTCAGCAGAGCGCAGATACCAAATACTGTCCTTC

TAGTGTAGCCGTAGTTAGGCCACCACTTCAAGAACTCTGTAGCACCGCCTACATACCTCGCT
CTGCTAATCCTGTTACCAGTGGCTGCTGCCAGTGGCGATAAGTCGTGTCTTACCGGGTTGG
ACTCAAGACGATAGTTACCGGATAAGGCGCAGCGGTCGGGCTGAACGGGGGGTTCGTGCA
CACAGCCCAGCTTGGAGCGAACGACCTACACCGAACTGAGATACCTACAGCGTGAGCTATG
AGAAAGCGCCACGCTTCCCGAAGGGAGAAAGGCGGACAGGTATCCGGTAAGCGGCAGGG
TCGGAACAGGAGAGCGCACGAGGGAGCTTCCAGGGGGAAACGCCTGGTATCTTTATAGTC
CTGTCGGGTTTCGCCACCTCTGACTTGAGCGTCGATTTTTGTGATGCTCGTCAGGGGGGC
GGAGCCTATGGAAAAACGCCAGCAACGCGGCCTTTTTACGGTTCCTGGCCTTTTGCTGGCC
TTTTGCTCACATGTTCTTTCCTGCGTTATCCCCTGATTCTGTGGATAACCGTATTACCGCCTT
TGAGTGAGCTGATACCGCTCGCCGCAGCCGAACGACCGAGCGCAGCGAGTCAGTGAGCG
AGGAAGCGGAAGAGCGCCTGATGCGGTATTTTCTCCTTACGCATCTGTGCGGTATTTCACA
CCGCAATGGTGCACTCTCAGTACAATCTGCTCTGATGCCGCATAGTTAAGCCAGTATACACT
CCGCTATCGCTACGTGACTGGGTCATGGCTGCGCCCCGACACCCGCCAACACCCGCTGAC
GCGCCCTGACGGGCTTGTCTGCTCCCGGCATCCGCTTACAGACAAGCTGTGACCGTCTCC
GGGAGCTGCATGTGTCAGAGGTTTTCACCGTCATCACCGAAACGCGCGAGGCAGCTGCGG
TAAAGCTCATCAGCGTGGTCGTGAAGCGATTCACAGATGTCTGCCTGTTCATCCGCGTCCA
GCTCGTTGAGTTTCTCCAGAAGCGTTAATGTCTGGCTTCTGATAAAGCGGGCCATGTTAAGG
GCGGTTTTTTCCTGTTTGGTCACTGATGCCTCCGTGTAAGGGGGATTTCTGTTCATGGGGG
TAATGATACCGATGAAACGAGAGAGGATGCTCACGATACGGGTTACTGATGATGAACATGCC
CGGTTACTGGAACGTTGTGAGGGTAAACAACTGGCGGTATGGATGCGGCGGGACCAGAGA
AAAATCACTCAGGGTCAATGCCAGCGCTTCGTTAATACAGATGTAGGTGTTCCACAGGGTAG
CCAGCAGCATCCTGCGATGCAGATCCGGAACATAATGGTGCAGGGCGCTGACTTCCGCGTT
TCCAGACTTTACGAAACACGGAAACCGAAGACCATTCATGTTGTTGCTCAGGTCGCAGACG
TTTTGCAGCAGCAGTCGCTTCACGTTCGCTCGCGTATCGGTGATTCATTCTGCTAACCAGTA
AGGCAACCCCGCCAGCCTAGCCGGGTCCTCAACGACAGGAGCACGATCATGCGCACCCGT
GGGGCCGCCATGCCGGCGATAATGGCCTGCTTCTCGCCGAAACGTTTGGTGGCGGGACCA
GTGACGAAGGCTTGAGCGAGGGCGTGCAAGATTCCGAATACCGCAAGCGACAGGCCGATC
ATCGTCGCGCTCCAGCGAAAGCGGTCCTCGCCGAAAATGACCCAGAGCGCTGCCGGCAC
CTGTCCTACGAGTTGCATGATAAAGAAGACAGTCATAAGTGCGGCGACGATAGTCATGCCCC
GCGCCCACCGGAAGGAGCTGACTGGGTTGAAGGCTCTCAAGGGCATCGGTCGAGATCCC
GGTGCCTAATGAGTGAGCTAACTTACATTAATTGCGTTGCGCTCACTGCCCGCTTTCCAGTC
GGGAAACCTGTCGTGCCAGCTGCATTAATGAATCGGCCAACGCGCGGGGAGAGGCGGTTT
GCGTATTGGGCGCCAGGGTGGTTTTTCTTTTCACCAGTGAGACGGGCAACAGCTGATTGCC
CTTCACCGCCTGGCCCTGAGAGAGTTGCAGCAAGCGGTCCACGCTGGTTTGCCCCAGCAG
GCGAAAATCCTGTTTGATGGTGGTTAACGGCGGGATATAACATGAGCTGTCTTCGGTATCGT
CGTATCCCACTACCGAGATATCCGCACCAACGCGCAGCCCGGACTCGGTAATGGCGCGCAT
TGCGCCCAGCGCCATCTGATCGTTGGCAACCAGCATCGCAGTGGGAACGATGCCCTCATTC
AGCATTTGCATGGTTTGTTGAAAACCGGACATGGCACTCCAGTCGCCTTCCCGTTCCGCTAT
CGGCTGAATTTGATTGCGAGTGAGATATTTATGCCAGCCAGCCAGACGCAGACGCGCCGAG
ACAGAACTTAATGGGCCCGCTAACAGCGCGATTTGCTGGTGACCCAATGCGACCAGATGCT
CCACGCCCAGTCGCGTACCGTCTTCATGGGAGAAAATAATACTGTTGATGGGTGTCTGGTC
AGAGACATCAAGAAATAACGCCGGAACATTAGTGCAGGCAGCTTCCACAGCAATGGCATCC
TGGTCATCCAGCGGATAGTTAATGATCAGCCCACTGACGCGTTGCGCGAGAAGATTGTGCA
CCGCCGCTTTACAGGCTTCGACGCCGCTTCGTTCTACCATCGACACCACCACGCTGGCAC

CCAGTTGATCGGCGCGAGATTTAATCGCCGCGACAATTTGCGACGGCGCGTGCAGGGCCA
GACTGGAGGTGGCAACGCCAATCAGCAACGACTGTTTGCCCGCCAGTTGTTGTGCCACGC
GGTTGGGAATGTAATTCAGCTCCGCCATCGCCGCTTCCACTTTTTCCCGCGTTTTCGCAGA
AACGTGGCTGGCCTGGTTCACCACGCGGGAAACGGTCTGATAAGAGACACCGGCATACTC
TGCGACATCGTATAACGTTACTGGTTTCACATTCACCACCCTGAATTGACTCTCTTCCGGGC
GCTATCATGCCATACCGCGAAAGGTTTTGCGCCATTCGATGGTGTCCGGGATCTCGACGCT
CTCCCTTATGCGACTCCTGCATTAGGAAGCAGCCCAGTAGTAGGTTGAGGCCGTTGAGCAC
CGCCGCCGCAAGGAATGGTGCATGCAAGGAGATGGCGCCCAACAGTCCCCCGGCCACGG
GGCCTGCCACCATACCCACGCCGAAACAAGCGCTCATGAGCCCGAAGTGGCGAGCCCGAT
CTTCCCCATCGGTGATGTCGGCGATATAGGCGCCAGCAACCGCACCTGTGGCGCCGGTGA
TGCCGGCCACGATGCGTCCGGCGTAGAGGATCGAGATCTCGATCCCGCGAAAT

**Protein sequence (CDS)**:
MGSSHHHHHHSSGLVPRGSHMDSIQAEEWYFGKITRRESERLLLNAENPRGTFLVRESETVKG
AYALSVSDFDNAKGLNVKHYLIRKLDSGGFYITSRTQFNSLQQLVAYYSKHADGLCHRLTTVCPT
SKMKPHPWFFGKIPRAKAEEMLSKQRHDGAFLIRESESVPGDFALSVKFGNDVQHFLVLRDGA
GKYFLWVVKFNSLNELVDYHRSTSVSRNQQIFLRDIEQVPQQP*

### Supporting Methods 1: Standard Operating Procedure for R2-pY

#### Standard Operation Procedure: Automated Phosphotyrosine Sample Preparation with R2-pY


1. Document Information 2
    1.1. Purpose 2
    1.2. Scope 2
    1.3. Outline 2
    1.4. Abbreviations 3
2. General Suggestions 3
3. R2-P1 (*optimized since Leutert et al. 2019*) 5
    3.1. Reagent and Material List 5
    3.2. Protocol 5-6
        3.2.1. Important Notes 6
        3.2.2. Experiment Planning 6
        3.2.3. Carboxylated Magnetic Bead Preparation 7
        3.2.4. Preparation of bead-ethanol mixture 7
        3.2.5. 96-well Plate Preparation for R2-P1 7-8
        3.2.6. R2-P1 KingFisher™ Flex Protocol 9-12
4. R2-pY 13
    4.1. Reagent and Material List 13
    4.2. Protocol 14
        4.2.1. Important Notes 14
        4.2.2. Experiment Planning 14-15
        4.2.3. Expression and purification of sSrc 15-17

4.2.4. sSrc Magnetic Bead Preparation 17-18

4.2.5. 96-well Plate Preparation for R2-pY 18-19

4.2.6. R2-pY KingFisher™ Flex Protocol 20-23

5. R2-P2 clean-up of R2-pY eluate 21

5.1. Reagent and Material List 21-22

5.2. Protocol 23

5.2.1. Important Notes 24

5.2.2. Experiment Planning 24

5.2.3. Magnetic $Fe^{3+}$-IMAC Bead Preparation and Recycling 25

5.2.4. 96-well Plate Preparation for R2-P2 25

5.2.5. R2-P2 KingFisher™ Flex Protocol 27-29

### 1. Document Information

#### a. Purpose

This standard operating protocol (SOP) document describes rapid-robotic-phosphotyrosine proteomics (R2pY) and rapid-robotic phosphoproteomics (R2-P2) protocols for automated sample processing to prepare phosphotyrosine (pY) proteomics samples in 96-well format for LC-MS analysis. R2-pY is a combination of pY-capture with SH2 superbinder or anti-pY antibodies conjugated to magnetic beads followed by phosphopeptide clean-up with $Fe^{3+}$-IMAC magnetic particles using a KingFisher™ Flex magnetic bead processor. We have tested various parameters of the method and provide a detailed protocol for the manual and automated steps.

#### b. Scope

The provided information will allow investigators to implement R2-pY as a standard sample processing method for a phosphotyrosine proteomic workflow resulting in MS-ready tyrosine phosphopeptide samples. Of note, R2-pY can be performed in addition to R2-P1 and R2-P2 described by Leutert et al. 2019 for parallel analysis of the total proteome and global phosphoproteome.

#### c. Outline

The document is organized in the following sections:

**General suggestions:** Considerations for planning experiments with R2-pY.

**Protocol for R2-P1:** A validated protocol for automated solid-phase-enhanced sample preparation in a 96-well plate format by a KingFisher™ Flex magnetic handling robot. Starts from cell or tissue lysates and ends with peptide samples ready for total proteome analysis by LC-MS and for phosphopeptide enrichments.

**Protocol for R2-pY:** A validated protocol for automated tyrosine phosphopeptide enrichment in a 96-well plate format by a KingFisher™ Flex magnetic handling robot. Starts from cell or tissue lysates and ends with peptide samples ready for stringent clean-up with R2-P2.

**Protocol for pY-peptide clean-up with R2-P2:** Combination of R2-pY and automated phosphopeptide enrichment carried out in a 96-well plate format by a KingFisher™ Flex magnetic handling robot. Starts with pY-enriched peptides and ends with samples ready for tyrosine phosphoproteome analysis by LC-MS.

#### d. Abbreviations

R2-P1 – rapid-robotic-proteomics
R2-pY – rapid-robotic phosphotyrosine proteomics
R2-P2 – rapid-robotic phosphoproteomics
sSrc – Src SH2 pY superbinder protein
NHS – *N*-hydroxy-succinimide functional group
p-Tyr-100 – Cell Signaling anti-phosphotyrosine antibody
p-Tyr-1000 – Cell Signaling anti-phosphotyrosine antibody
Ab – antibody
CST - Cell Signaling Technology ®
ACN – acetonitrile
AmBic – ammonium bicarbonate
AP – Affinity Purifcation (20 mM MOPS-NaOH, 10 mM $Na_2HPO_4$, NaCl, pH 7.2)
EtOH – ethanol
IMAC – Immobilized-Metal Affinity Chromatography
LC – Liquid Chromatography
MS – Mass Spectrometry
BCA - Bicinchoninic acid
MeOH – Methanol
TFA – Trifluoroacetic Acid
IPTG - Isopropyl β-d-1-thiogalactopyranoside
PBS - phosphate buffered saline
NaCl - sodium chloride
SDS PAGE - Sodium dodecyl sulfate polyacrylamide gel electrophoresis

### 2. General Suggestions

**Protease inhibitors:** As outlined in Leutert *et al.* 2019, Pierce protease inhibitors (#A32963) are not efficiently removed by R2-P1 and lead to decreased digestion efficiency by trypsin or LysC if they are present at >0.2x in the digestion buffer. To circumvent this, one of the following strategies, or a combination of them, can be chosen:

- Use a denaturing buffer (e.g. 8 M urea) without protease inhibitors for lysis.
- Produce cell lysates at protein concentrations higher than 4 mg/mL.
- Dilute lysates 4-fold prior to digestion.

**Phosphatase inhibitors:** As outlined in Leutert *et al.* 2019, a common phosphatase inhibitor mix (50 mM sodium fluoride, 10 mM sodium pyrophosphate, 50 mM sodium beta-glycerophosphate, 1 mM sodium orthovanadate) is not efficiently removed by R2-P1 and competes during phosphopeptide enrichment. Therefore, we recommend using denaturing buffers (e.g. 8 M urea) without phosphatase inhibitors for lysis ahead of R2-pY and R2-P2.

**Precipitates:** Depending on the sample, we have observed insoluble particles due to precipitation at a few steps in the protocol that we indicate below. It is critical that tyrosine phosphopeptide and global phosphopeptide enrichments are performed on a clarified peptide mixture for maximal efficiency. If precipitation is observed at any step after protein digestion the plate needs to be centrifuged for 10 minutes at maximal speed and supernatant transferred to a new plate. If this does not help to remove visible precipitates, longer centrifugation is needed. Alternatively, samples can be transferred to PCR tube strips or microtubes and centrifuged at higher speeds.

**Magnetic particles for phosphopeptide enrichment:** The phosphopeptide clean-up described in this protocol is performed using $Fe^{3+}$-IMAC magnetic particles. As outlined by Leutert et al. 2019, $Fe^{3+}$-IMAC is the most efficient material for phosphopeptide enrichment; it is easy to use and cheap (since the $Fe^{3+}$ can be stripped and reloaded multiple times). However, phosphopeptide clean-up may also work with $Ti^{4+}$-IMAC, $TiO_2$, $Zr^{4+}$-IMAC microspheres from MagReSyn following the provided instructions. We have not evaluated the alternative resins for clean-up of pY-peptides, however, the resins were of comparable quality during validation of R2-P2. However, caution should be taken as it is possible that alternative R2-P2 resins provide slightly distinct phosphopeptide selectivity as described by Leutert *et al.* 2019.

**Expression and Purification of sSrc:** We recommend preparing purified sSrc in advance. Each *Escherichia coli* cell pellet and purified sSrc can be stored at -80°C for at least one year. sSrc was expressed in NEB T7 express cells, however similar cells may work as well.
We have found sSrc in phosphate buffered saline with 0.05% sodium azide is stable through freeze thaw. However, other buffers may be suitable as well.

# 3. R2-P1

#### a. Reagent and Material List

| Item | Vendor | Catalog # |
|---|---|---|
| KingFisher™ Flex | Thermo Fisher Scientific | |
| KingFisher 96 KF microplate (200 µL) | Thermo Fisher Scientific | 97002540 |
| KingFisher 96 microtiter DW plate | Thermo Fisher Scientific | 95040450 |
| KingFisher 96 tip comb for DW magnets | Thermo Fisher Scientific | 97002534 |
| SpeedVac Vacuum Concentrator | various | |
| Plate, PCR or microtube centrifuge | various | |
| Sera-Mag SpeedBead Carboxylate-Modified Magnetic Particles (Hydrophilic) | Cytiva Life Sciences | 45152105050250 |
| Sera-Mag SpeedBead Carboxylate-Modified Magnetic Particles (Hydrophobic) | Cytiva Life Sciences | 65152105050250 |
| Water LC-MS grade | Fisher Scientific | 7732-18-5 |
| Acetonitrile LC-MS grade | Fisher Scientific | 75-05-8 |
| Formic acid LC-MS grade | Fisher Scientific | A117-50 |
| Ethanol 200 Proof (100%) | Decon Labs | 2701 |
| Ammonium bicarbonate | Sigma-Aldrich | A6141 |
| Sequencing grade modified trypsin | Promega | V5111 |
| Methanol LC-MS grade (optional) | Fisher Scientific | A456-4 |
| Acetic acid glacial HPLC grade (optional) | Fisher Scientific | A35-500 |
| C8 Extraction Disks 3M™ Empore™ (optional) | Fisher Scientific | 14-386 |

#### b. Protocol

##### i. Important Notes

1. All steps are performed at room temperature, unless stated otherwise.
2. The following protocol is for 1000 µg of protein digest in one well. The protocol has been successfully tested for protein input amounts from 25 µg to 1000 µg.

3. The protocol has been extensively tested with lysates performed in urea buffer (8 M urea, 150 mM NaCl, 100 mM Tris pH 8), however other lysis buffers should work as well.
4. With the recommended buffer volumes and ratios, the maximal amount of protein that can be processed in one well is 1000 µg.
5. We perform digests for 4 hours at 37 ºC, however these parameters can easily be adjusted as needed in the protocol.
6. All plate pipetting steps are performed with 8- or 12- multi-channel pipettes.

#### ii. Experiment planning

1. Determine the amount of protein sample that needs to be processed, considering:
    a. For proteome analysis only, we recommend starting with 25 µg of protein.
    b. For phosphoproteomic analysis we recommend starting with a minimum of 100 µg protein, with an optimal starting amount of 200 µg – 500 µg. This is sample dependent and needs to be tested.
2. Determine the protein concentration of clarified cell lysate(s) by BCA assay or equivalent. Reduce and alkylate cysteines by preferred method. Adjust protein concentration in samples to 5 µg/µl with lysis buffer.
3. Determine the amount of magnetic carboxylated beads to be used. We recommend using 1 µl of 10 µg/µl of carboxylated bead mix per µg of protein to be processed; e.g. 10 mg of beads per 1 mg of protein input.
4. Determine the volumes of the solutions to be used per well in the different plates:
    a. Binding plate:
        1. Lysate volume ($V_{Lysate}$) = Calculate lysate volume to reach desired protein amount, keeping a protein concentration of 5 µg/µl.
        2. Water volume = ~0.1 mL per 10 mg beads
        3. 200 Proof Ethanol volume = 0.7 - 0.8 mL per well.
    b. Wash plates (4x): 80% EtOH volume = 0.8 mL per well
    c. Elution 1 plate:
        i. 500 µl 25 mM AmBic pH 8.2
            1. Less volume may for less protein input. Acidification buffer should scale as well.
        ii. Trypsin or LysC: 1 µg enzyme : 100 µg protein
    d. Elution 2 plate: 100 µl water
5. Determine the type of plates to be used for the different solutions: For volumes of 50 µl – 150 µl per well use Kingfisher microplates

(shallow well) and for 150 µl – 1000 µl per well use Kingfisher microtiter DW plates (deep well).

##### iii. Carboxylated magnetic bead preparation

1. Take both (hydrophilic and hydrophobic) magnetic carboxylated bead stocks off the fridge, warm to room temperature and vortex gently to fully suspend magnetic beads.
2. Beads come at a stock concentration of 50 µg/µl. Take the required amount of beads and mix hydrophilic and hydrophobic beads at a 1:1 ratio. Dilute to a total bead concentration of 1 µg/µl, wash the beads three times with water keeping the beads at 1 µg/µl on a magnetic rack in an Eppendorf or Falcon tube and resuspend beads in water at the working concentration of 10 µg/µl.

##### iv. Preparation of bead-ethanol mixture

Beads will be resuspended in 70% ethanol immediately before plate preparation. It is important to keep beads suspended in a small amount of water before addition of ethanol to avoid clumps.

1. Resuspend the required amount of beads at 100 mg/mL in water. This means 10 mg of beads will be resuspended to a total volume of 0.1 mL in water.
2. Add 200 Proof ethanol to reach a total volume of 800 µL. For example, 10 mg of beads resuspended in ethanol to 0.8 mL will yield a 12.5 µg/µL bead solution in 70% ethanol. Samples digesting less than 1 mg of protein input will yield ethanol concentrations closer to 80% final.
3. This bead-ethanol solution is now ready for addition to lysate during plate preparation.

##### v. 96-well plate preparation for R2-P1

1. Prepare an empty plate containing the tip comb.
2. Prepare a binding plate in the following way:
   a. Add the calculated volumes of lysate to each well. If lysate volume is below 200 µL, fill up to 200 µL with lysis buffer.
   b. Add the required volume of bead-ethanol mixture into the lysate in the wells. No mixing or pipetting required.
      i. e.g.: Wells with 1 mg of protein input will receive 0.8 mL containing 10 mg of beads in 70% ethanol.
3. Prepare the 4 Wash plates by dispensing 80% EtOH.
4. Prepare an Elution-2 plate by dispensing water.
5. Start the R2-P1 Kingfisher program and follow the instructions on the robot. Leave the space for the Elution-1 plate empty and start the protocol.
6. Start preparing Elution-1 plate on ice 25 min into the protocol.

7. The R2-P1 Kingfisher program will pause after approximately 35 minutes. Follow the instructions of the robot, load Elution-1 plate and resume the program.
8. Remove all plates after the program finishes.
9. Transfer solution in Elution-2 plate to Elution-1 plate.
10. Stop the enzymatic digestion by adding formic acid to the digest to pH < 2 (usually final formic acid concentration is 1-5%).
    a. For example, 60 µL of 10% formic acid, 75% ACN may be used to acidify the 600 µL of pooled elutions.
    b. For total proteome measurement, pH adjustment is particularly critical.
11. Depending on the sample, varying degrees of precipitation might be observed after digestion. For efficient downstream sample processing (e.g. phosphopeptide enrichment or LC-MS analysis) remove precipitates by centrifugation and transfer the supernatant to a new plate.
12. Take a sample aliquot for total proteome analysis from supernatant, see optional filtering step below or transfer to a MS vial or MS sample plate.
13. (Optional step that we recommended when processing many samples in parallel): To avoid any carryover of magnetic beads to the LC system, filter samples through a C8 stage tip or stage tip plate. For this, pack a stage tip with one layer of C8 material and perform the following steps:
    a. Condition with 30 µl MeOH.
    b. Wash with 30 µl 100% ACN.
    c. Wash with 30 µl 70% ACN, 0.25% acetic acid.
    d. Hold stage tip on top of MS sample vial.
    e. Adjust sample to 50% ACN and pass sample through stage tip, collecting eluate in the MS vial.
    f. Elute with 30 µl 70% ACN, 0.25% acetic acid, collecting this second eluate also in the MS vial.
14. Dry samples down in a speedvac. Dried samples can be stored at -20 ºC. Resuspend in 4% formic acid and 3% ACN prior to LC-MS analysis.
15. The rest of the plate is dried down and can be stored at -20 ºC for subsequent phosphopeptide enrichment with R2-pY to R2-P2 for phosphotyrosine-specific enrichment or R2-P2 alone for global phosphopeptide enrichment.

### vi. R2-P1 KingFisher™ Flex protocol



#### General info

##### Protocol information

| | |
|---|---|
| Protocol name | R2-P1_premix_5h_2022_1mg_input |
| Modified by | KingFisher |
| Kit name | R2-P1 |
| Description | R2-P1 for desalting and digestion of 1 mg of protein per well.<br>[Alexis Chang, Villen Laboratory, University of Washington]<br><br>!!!!Beads and Protein Mixed!!!!<br><br>!!!4.3h digestion!!! |



#### Sample layout

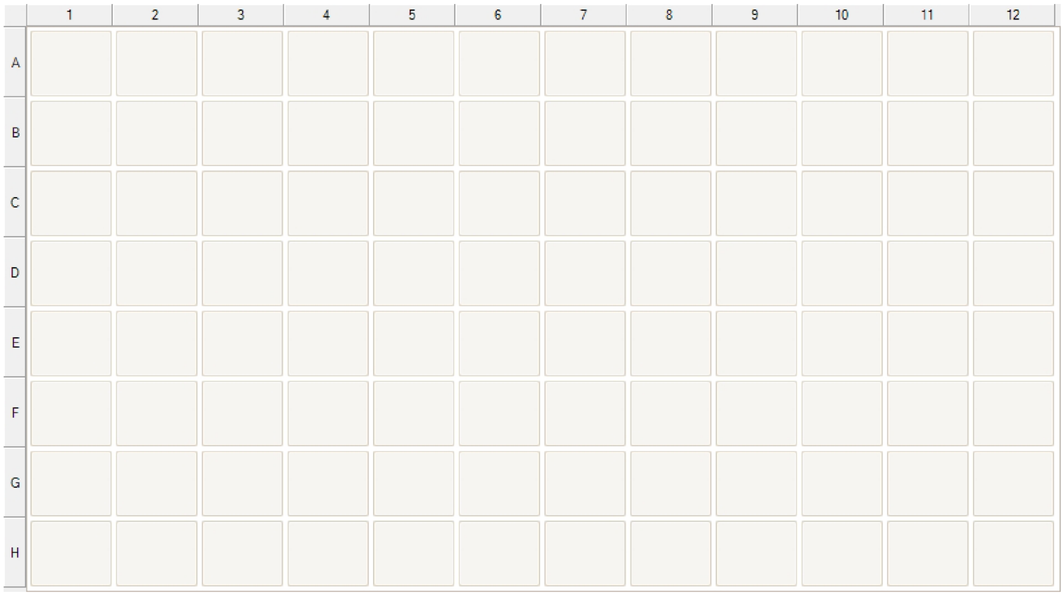

#### Reagent info

| Wash 4 | | 96 DW plate | |
|---|---|---|---|
| Name | Well volume [µl] | Total reagent volume [µl] | Type |
| 80% EtOH | 800 | - | Reagent |

| Binding | | 96 DW plate | |
|---|---|---|---|
| Name | Well volume [µl] | Total reagent volume [µl] | Type |
| 100% EtOH | 550 | - | Sample |
| Lysate (5ug/ul protein) | 200 | - | Reagent |
| 1x beads in 99.9% EtOH | 250 | - | Reagent |

| Wash 1 | | 96 DW plate | |
|---|---|---|---|
| Name | Well volume [µl] | Total reagent volume [µl] | Type |
| 80% EtOH | 800 | - | Reagent |

| Wash 2 | | 96 DW plate | |
|---|---|---|---|
| Name | Well volume [µl] | Total reagent volume [µl] | Type |
| 80% EtOH | 800 | - | Reagent |

| Wash 3 | | 96 DW plate | |
|---|---|---|---|
| Name | Well volume [µl] | Total reagent volume [µl] | Type |
| 80% EtOH | 800 | - | Reagent |

| Tip Storage Plate | | 96 standard plate | |
|---|---|---|---|
| Name | Well volume [µl] | Total reagent volume [µl] | Type |
| - | - | - | - |

| Elution 1 - Trypsin | | 96 DW plate | |
|---|---|---|---|
| Name | Well volume [µl] | Total reagent volume [µl] | Type |
| 25mM Ambic, Trypsin 1:100 | 500 | - | Reagent |

| Elution 2 | | 96 standard plate | |
|---|---|---|---|
| Name | Well volume [µl] | Total reagent volume [µl] | Type |
| H2O | 100 | - | Reagent |

#### Dispensed reagents

The protocol does not contain dispensed reagents

#### Steps data

| | | |
|---|---|---|
| **Tip1** | **96 DW tip comb** | |
| **Pick-Up** | **Tip Storage Plate** | |
| **Capture Proteins** | **Binding** | |
| Beginning of step | Precollect | No |
| | Release beads | No |
| Mixing / heating: | Mixing time, speed | 00:13:00, Slow |
| | Heating during mixing | No |
| End of step | Postmix | No |
| | Collect count | 5 |
| | Collect time [s] | 30 |
| **Wash 1** | **Wash 1** | |
| Beginning of step | Precollect | No |
| | Release time, speed | 00:00:30, Fast |
| Mixing / heating: | Mixing time, speed | 00:02:00, Slow |
| | Heating during mixing | No |
| End of step | Postmix | No |
| | Collect count | 5 |
| | Collect time [s] | 30 |
| **Wash2** | **Wash 2** | |
| Beginning of step | Precollect | No |
| | Release time, speed | 00:00:30, Fast |
| Mixing / heating: | Mixing time, speed | 00:02:00, Slow |
| | Heating during mixing | No |
| End of step | Postmix | No |
| | Collect count | 5 |
| | Collect time [s] | 30 |
| **Wash3** | **Wash 3** | |
| Beginning of step | Precollect | No |
| | Release time, speed | 00:00:30, Fast |
| Mixing / heating: | Mixing time, speed | 00:02:00, Slow |
| | Heating during mixing | No |
| End of step | Postmix | No |
| | Collect count | 5 |
| | Collect time [s] | 30 |
| **Wash4** | **Wash 4** | |
| Beginning of step | Precollect | No |
| | Release time, speed | 00:00:30, Slow |
| Mixing / heating: | Shake 1 time, speed | 00:02:00, Slow |
| | Shake 2 time, speed | 00:00:30, Fast |
| | Heating during mixing | No |
| End of step | Postmix | No |
| | Collect beads | No |

| | | | |
|---|---|---|---|
| Pause - Load Elution1 | | Elution 1 - Trypsin | |
| | | Message | Insert Elution1 - Trypsin |
| | | Dispensing volume [µl] | 0 |
| Pause - Load Elution2 | | Elution 2 | |
| | | Message | Insert Elution2 |
| | | Dispensing volume [µl] | 0 |
| Collect Beads Wash 4 | | Wash 4 | |
| | | Collect count | 5 |
| | | Collect time [s] | 30 |
| Protein Digestion and Elution | | Elution 1 - Trypsin | |
| | Beginning of step | Precollect | No |
| | | Release time, speed | 00:01:00, Bottom mix |
| | Mixing / heating: | Shake 1 time, speed | 00:00:15, Bottom mix |
| | | Shake 2 time, speed | 00:00:15, Fast |
| | | Shake 3 time, speed | 00:04:30, Slow |
| | | Loop count | 52 |
| | | Heating temperature [°C] | 37 |
| | | Preheat | No |
| | End of step | Postmix | No |
| | | Collect count | 5 |
| | | Collect time [s] | 30 |
| Elution 2 | | Elution 2 | |
| | Beginning of step | Precollect | No |
| | | Release time, speed | 00:01:00, Bottom mix |
| | Mixing / heating: | Shake 1 time, speed | 00:01:00, Bottom mix |
| | | Shake 2 time, speed | 00:04:00, Medium |
| | | Heating during mixing | No |
| | End of step | Postmix | No |
| | | Collect count | 5 |
| | | Collect time [s] | 30 |
| Dispose Beads | | Wash 4 | |
| | | Release time, speed | 00:00:30, Bottom mix |
| Leave | | Tip Storage Plate | |

#### Lot info

No lot numbers have been defined.

# 4. R2-pY

#### a. Reagent and Material List

| Item | Vendor | Catalog # |
|---|---|---|
| KingFisher™ Flex | Thermo Fisher Scientific | |
| KingFisher 96 KF microplate (200 µL) | Thermo Fisher Scientific | 97002540 |
| KingFisher 96 microtiter DW plate | Thermo Fisher Scientific | 95040450 |
| KingFisher 96 tip comb for DW magnets | Thermo Fisher Scientific | 97002534 |
| SpeedVac Vacuum Concentrator | various | |
| Plate, PCR or microtube centrifuge | various | |
| Bath sonicator | various | |
| Pierce™ NHS-Activated Magnetic Beads | Thermo Scientific | 88826 |
| p-Tyr-100 magnetic bead conjugate | Cell Signaling Technology | 8095S |
| p-Tyr-1000 magnetic bead conjugate | Cell Signaling Technology | 14017S |
| T7 Express Competent E. *coli* cells | New England Biolabs | C2566H |
| Water LC-MS grade | Fisher Scientific | W6500 |
| Isopropyl β-D-1-thiogalactopyranoside | Gold Bio | 12481C25 |
| Phosphate buffered saline tablets | Fisher Scientific | BP2944-100 |
| TALON® Metal Affinity Resin | Takara Bio | 635503 |
| Pierce™ 660 nm Protein Assay Reagent | Thermo Fisher Scientific | 22660 |
| Pierce™ BCA Protein Assay Kit | Thermo Fisher Scientific | 23227 |
| Acetonitrile LC-MS grade | Fisher Scientific | 75-05-8 |
| Formic acid LC-MS grade | Fisher Scientific | A117-50 |
| Trifluoroacetic acid LC-MS grade | Fisher Scientific | A116-50 |
| MOPS free acid | Fisher Scientific | BP308-100 |
| Sodium acetate | Sigma-Aldrich | S2889-250G |
| Sodium phosphate monobasic | Sigma-Aldrich | S0751-500G |
| Sodium chloride | Sigma-Aldrich | S3014-500G |
| Sodium hydroxide | Fisher Scientific | S318-500 |

#### b. Protocol

##### i. Important Notes

- All steps are performed at room temperature unless stated otherwise.
- The following protocol is for 1 mg of protein digest in one well. The protocol has been successfully tested for protein input amounts from 250 µg to 4 mg.
- The protocol has been extensively tested with peptides in 1X AP (20 mM MOPS-NaOH, 10 mM $Na_2HPO_4$, NaCl, pH 7.2), however other peptide resuspension and wash buffers may work as well.
- With the recommended bead size and slurry concentration, the maximum amount of Pierce™ NHS magnetic beads per well is 1.8 mg. If more than 1.5 mg of beads are used, bead slurry must be concentrated beyond 10 µg/µL to reach a maximum volume of 150 µL to fit within a shallow 96-well plate.
- We recommend the maximum amount of protein that can be processed in one well is 4 mg. If more than 4 mg of starting protein is desired, we recommend splitting the sample across multiple wells and combining final peptide samples prior to MS measurement.
- We perform incubation of peptides and anti-pY beads for 2 hours. However, it is possible less than 2 hours is sufficient interaction time for capture of pY-peptides.
- All plate pipetting steps are performed with 8- or 12- multi-channel pipettes.

##### ii. Experiment Planning

- Determine the amount of protein sample that needs to be processed, considering for tyrosine phosphoproteomic analysis we recommend starting with a minimum of 1 mg of protein input.
    - This is sample dependent and needs to be tested.
    - Samples with low amounts of pY modifications will require more protein input while samples with higher levels of pY modifications will require less protein input.
- Determine the protein concentration of clarified cell lysate(s) by BCA assay or equivalent. Reduce and alkylate cysteines by preferred method. Digest and desalt peptide by preferred method:
    - Options include: R2-P1 or solution digest with cartridge desalting.
- Dry peptides by vacuum centrifugation and resuspend peptides in 1X AP pH 7.2 such that each well receives 900 µL of peptide suspension.
    - Ensure pH of peptide suspension is between 7 to 8.
- Determine the amount of magnetic pY-capture beads to be used. We recommend using 80 µL - 100 µL of NHS-conjugted sSrc magnetic beads (10 µg/µL) or CST® antibody beads (proprietary slurry concentration) per 0.5 mg - 2 mg of sample protein.
    - Importantly, to compare pY enrichment results between samples, the bead to peptide ratio should be kept constant (Fig. 4) .
- Determine the volumes of the solutions to be used per well in the different plates:
    - Binding plate:
        - 900 uL of 1X AP buffer will be needed per well.

- Wash plates 1 - 3 (3X):
  - 1 mL of 1X AP buffer will be needed per well.
- Wash plate 4:
  - 1 mL of hplc-grade water will be needed per well.
- Elution plate 1:
  - 100 µL of 0.5% TFA will be needed per well.
- Elution plate 2:
  - 150 µL of 1% TFA, 60% ACN will be needed per well.

- Determine the type of plates to be used for the different solutions: For volumes of 50 – 150 µL per well use KingFisher microplates (shallow well) and for 150 µL – 1000 µL per well use KingFisher microtiter DW plates (deep well).

##### iii. Expression and purification of sSrc

- Determine the amount of sSrc desired. Roughly, you will need a 1 L culture of E. *coli* expressing sSrc per 96-well plate.
  - This ratio is based on the observed yield of 20 mg of TALON®-purified sSrc per liter of culture. The ratio may vary due to slight variations in culture conditions, purification efficiency, and sample handling.
  - *Example calculations:* If 80 µL of NHS-sSrc beads are used per well, you will need 80 µL of sSrc at 1.5 - 2 mg/mL for bead conjugation. Therefore, each well will require a total mass of 120 µg - 160 µg of purified sSrc. If all 96-wells are used, 11.5 - 15.4 mgs of sSrc will be needed for conjugation. Therefore, we recommend scaling predictions of sSrc amount requirements on the number of wells to be used. Still, it is advised to grow and purify excess sSrc. Unused cell pellets or purified sSrc in PBS pH 7.4 (pH 7-8) can be stored at -80 °C for future conjugations.
- Culturing
  - Inoculate small volume cultures from frozen glycerol stocks of NEB T7 express containing sequence-verified pET28a(+)_sSrc using appropriate antibiotic (i.e. 50 mg/mL kanamycin). Culture volume should allow for inoculation of larger culture with 100-fold dilution.
  - Grow inoculation cultures for 12 - 18 hours.
  - Inoculate each flask of sterile Luria broth with inoculation culture at 1:50 to 1:100 dilution.
  - Incubate at 36 - 38°C until optical density at 600 nm ($OD_{600}$) reaches 0.3 to 0.4.
  - Incubate cultures at 18 - 20°C until $OD_{600}$ reaches 0.6 - 0.8 then induce with 1 mM IPTG.
  - Grow induced cultures at 18 - 20°C for 20 - 24 hrs.
  - Collect cells by centrifugation and discard media. Always store cells on ice or chilled between handling.
  - Either proceed immediately to cell lysis and purification or store cell pellets at -80°C.
- Cell lysis
  - At all times during lysis, keep samples on ice or chilled.

    - Resuspend cell pellet in lysis buffer (1X PBS, 0.1 mg/mL lysozyme, 12 U/mL benzonase nuclease, pH 7.4) at approximately 5 - 6 mL lysis buffer per gram of cell pellet.
    - Incubate cell suspension for 30 minutes at 4°C with gentle mixing to allow lysozyme to soften cell walls and benzonase to degrade nucleic acids.
    - Sonicate cell suspension 6 times each for 30 - 45 seconds with 45 - 60 second intermittent rests all on ice. Aim for moderate sonication power, such as 40-45% or 12 watts. Avoid sample overheating or frothing.
    - Incubate lysate at 4 - 8°C for 10 min with gentle mixing to allow further degradation of nucleic acids.
    - Clarify lysate by chilled centrifugation such as 16,000 - 20,000 x g for 30 minutes at 4°C. Collect soluble supernatant and discard insoluble pellets.
        - Optional: further clarify lysate by 0.45 µm syringe filtration.
        - Save 20 µL of clarified lysate for SDS PAGE analysis.
- Purification of sSrc with TALON® (Cobalt 2+) resin
    - sSrc contains a six-histidine affinity tag on its N-terminus to enable purification ahead of conjugation to magnetic beads. Purification of sSrc may be performed in a number of ways.
        - For example, cobalt(II) or nickel-NTA affinity should work. We tested cobalt affinity for its weaker interaction and therefore more stringent purification of sSrc.
        - Purification can be performed in batch mode, by gravity or on an FPLC. We tested batch and gravity purification and achieved good results.
            - Within batch and gravity purification with TALON® resin, we found a ratio of 2 mL dry resin volume (or 4 mL of 50% slurry) saturated capture of sSrc from approximately 5 g of cell pellet or 1 L culture. Still, the ideal ratio of resin to cell amount may be broader.
        - Further, we used a 1X phosphate saline buffer and eluted with imidazole, however other buffer and elution conditions may also work.
    - Setup a gravity or batch column with 2 mL of dry resin volume (4 mL of 50% slurry) per 1 L culture or ~ 5 g cells.
    - Equilibrate TALON resin in 1X PBS pH 7.4
    - Combine equilibrated resin with clarified lysate and incubate with gentle mixing for 30 - 60 min at 4 - 8°C.
    - Collect flow through.
        - Optional: Save 20 µL of clarified lysate for SDS PAGE analysis.
    - Wash column twice with 5 - 10 column volumes (CVs) of 1X PBS pH 7.4.
    - Wash column once with 5- 10 CVs of 1X PBS, + additional 150 mM NaCl, pH 7.4.
    - Wash column once with 2 CVs of 1X PBS, 5 mM imidazole, pH 7.4
    - Wash column once with 1 CVs of 1X PBS, 10 mM imidazole, pH 7.4
        - Note: significant amounts of sSrc will elute here along with contaminants.
    - Wash column twice with 2 CVs of 1X PBS pH 7.4.
    - Into fresh vessels, collect purified sSrc with six to eight 1CV fractions of 1X PBS, 300 mM imidazole, pH 7.0 - 7.4.

- Recommended: Allow each addition of elution buffer to incubate with column for 2 min prior to collection.

- Regeneration and storage of TALON resin:

Wash resin with 20 mM 2-(N-morpholine)-ethanesulfonic acid (MES) buffer (pH 5.0) before reuse to remove bound imidazole. Note: do not store TALON in denaturants.

  - Wash resin with 5 CVs of 20 mM MES, 0.3 M NaCl, pH 5.0.
  - Wash resin with 5 CVs of distilled water.
  - Store resin at 4 - 8°C in 20% non-buffered ethanol, 0.1% sodium azide.
- Buffer exchange
  - Run an SDS PAGE to determine which sSrc elution fractions to pool. Pool elution fractions that display a reasonably pure band near 15 kDa. True molecular weight of the sSrc construct is 14.7 kDa however we observed migration at slightly higher molecular weight, around 15 - 16 kDa (Supporting Figure 1). For example, we pooled all eight elution fractions, seven of which are displayed in the SDS PAGE in Supporting Figure 1 (lanes 8 - 14).
  - Other fractions such as load, flow through, washes may be run if desired. Examples of load, flow-through and washes are shown in Supporting Figure 1.
- If proceeding to conjugation, exchange pooled sSrc into 50 mM borate pH 8.5 and concentrate to 1.5 to 2 mg/mL sSrc concentration.
  - Buffer exchange is commonly achieved with dialysis or spin concentration. The goal is to remove residual imidazole and transfer sSrc into the conjugation buffer.
- Alternatively, if storing sSrc at -80°C for later use, exchange pooled sSrc into 1X PBS pH 7.4 such as with dialysis or spin concentration. The goal is to remove residual imidazole.
- Quantify sSrc with appropriate method (i.e. BCA assay, absorbance at 280 nm, Bradford assay or 660 nm assay, among others).

**sSrc protein parameters**

| construct | Molecular weight (Da) | Mass Extinction coefficient | Theoretical Isoelectric point |
|---|---|---|---|
| **His$_6$ - thrombin cut site - sSrc$^{T8V, C10A, K15L}$**<br><br>MGSS-**HHHHHH**-SSG-**LVPRGS**-HM-**DSIQAEE WYFGKITRRESERLLLNAENPRGTFLVRESET VKGAYALSVSDFDNAKGLNVKHYLIRKLDSGG FYITSRTQFNSLQQLVAYYSKHADGLCHRLTTV CPTSK** | **14716.51 Da** | **0.981** | **9.20** |

#### iv. sSrc Magnetic Bead Preparation

Allow at least one day to conjugate sSrc to magnetic beads before R2-pY. We have tested storage of sSrc-NHS beads at 4°C up to 1 week ahead of R2-pY. sSrc conjugated to magnetic beads are likely to tolerate longer storage times at 4°C however we did not test longer storage.

Magnetic bead preparation steps are performed on a magnetic rack in 1.5 - 2ml tubes. Ensure homogenous bead suspension during each washing step. To resuspend beads, we suggest tube inversion or gentle tapping. Following each wash, remove as much liquid as possible without losing beads.

- **Prepare:**
  - Prepare ice-cold 1 mM hydrochloric acid ahead of time.
  - Incubate both purified sSrc in 50 mM borate pH 8.5 and Pierce™ NHS-activated magnetic beads at room temperature for 30 min. Gently vortex the beads and tap to resuspend.
  - Aliquot bead slurry (10 µg/µL) across 1.5 to 2 mL microtubes in 200 to 400 µL aliquots.
- **Condition:** Remove bead storage solution. Add 1 mL of 1 mM HCl to each bead aliquot. Mix by gentle vortexing for 5-10 sec. Discard HCl wash.
  - Note: Beads fail to form a compact pellet after HCl wash. Therefore, it may not be possible to remove all HCl.
- **Bind:** Immediately add purified sSrc to each bead aliquot. Volume of purified sSrc will be equal to volume of 10 µg/µL bead slurry. For example, a 300 µL aliquot of beads requires 300 µL of sSrc at 1.5 to 2 mg/mL. Resuspend beads by gentle and brief tapping and vortexing.
  - Optional: Plan to retain a small amount of purified sSrc to serve as a baseline in quantification of conjugation efficiency. Conjugation efficiency can be measured by a 660 nm assay and may be qualitatively checked with SDS PAGE.
  - Incubate beads and protein for 2 hours at room temperature with inversion such that beads don't settle. Briefly vortex beads every 5 min for the first 30 min then every 15 min for the remaining 2 hrs.
  - Optional: Save the flow through for measurement of conjugation efficiency using a 660 nm assay and visual check with SDS PAGE.
- **Quench & Wash:** Wash beads three times each with 1 mL of 50 mM Tris, 100 mM NaCl, pH 8.0 - 8.1. Incubate beads with the third wash for 2 hrs at room temperature with inversion.
- **Store:** Wash beads three times with 50 mM borate, 150 mM NaCl, pH 8.5, 0.05% azide then store in the same buffer at 4°C.
  - Recommended: store beads less dilute than 10 µg/µL, such as 2- 5 µg/µL.

##### v. 96-well Plate Preparation for R2-pY

- **Tip & Bead plate (shallow or deep well, depends on bead volume):**
  - Prepare the sSrc-NHS magnetic beads by washing three times in 1X AP pH 7.2 where the volume of the last wash should leave beads at 10 µg/µL.
  - Add the calculated amount of sSrc beads to each well.
    - If more than 100 µL of bead slurry is used, switch to a deep well plate.
    - If different volumes of beads are used equalize volumes by addition of 1X AP pH 7.2.
- **Wash plates (deep well):**
  - Prepare 3 wash plates by dispensing 1 mL of 1X AP pH 7.2 to each well.
  - Prepare 1 wash plate by dispensing 1 mL of HPLC grade water to each well.
- **Elution plate 1 (shallow well):**

    - Dispense 100 µL of 0.5% TFA to each well.
- **Elution plate 2 (shallow well)**:
    - Do not fill until 130 min into the R2-pY protocol.
    - After 130 min into the R2-pY program, dispense 150 µL of 1% TFA, 60% ACN in HPLC grade water.
- Start the R2-pY program.
    - Leave the space for elution plate 2 empty.
    - Start preparing elution plate 2 after 130 minutes into the program.
- The R2-pY protocol will pause after approximately 140 min into the program. Follow the instructions of the robot, load elution plate 2 and resume the program.
- Remove all plates after the program finishes.
- Combine elution 1 and elution 2 either into microtubes or another deep well plate.
- Add 650 µL of 100% ACN to each pooled eluate to reach 80% ACN.
- Centrifuge pooled and equilibrated lysate at max speed for 10 minutes.
- Transfer supernatant to fresh wells within a 96-deep well plate.
- Proceed to R2-P2 protocol. Cover plates until further use.
- Alternatively, if R2-P2 must occur more than a few hours later, it may be possible to dry peptides by vacuum centrifugation, cover and store at -80°C.

### vi. R2-pY KingFisher™ Flex Protocol



#### General info

Protocol information

| | |
|---|---|
| Protocol name | pY_SuperBinder_v2 |
| Modified by | KingFisher |
| Kit name | p-Tyr peptide enrichment |
| Description | R2-pY: tyrosine phosphopeptide enrichment protocol. Either sSrc or antibody magnetic beads can be used.<br>[Alexis Chang, Villen Laboratory, University of Washinton] |

### Reagent info

| Peptides | | 96 DW plate | |
|---|---|---|---|
| Name | Well volume [µl] | Total reagent volume [µl] | Type |
| anti-pY beads and peptides in IAP | 1000 | - | Reagent |

| Wash1 | | 96 DW plate | |
|---|---|---|---|
| Name | Well volume [µl] | Total reagent volume [µl] | Type |
| 1x IAP | 1000 | - | Sample |

| Wash2 | | 96 DW plate | |
|---|---|---|---|
| Name | Well volume [µl] | Total reagent volume [µl] | Type |
| 1x IAP | 1000 | - | Reagent |

| Wash3 | | 96 DW plate | |
|---|---|---|---|
| Name | Well volume [µl] | Total reagent volume [µl] | Type |
| 1x IAP | 1000 | - | Reagent |

| H20 Wash | | 96 DW plate | |
|---|---|---|---|
| Name | Well volume [µl] | Total reagent volume [µl] | Type |
| H2O hplc grade | 1000 | - | Reagent |

| Elution 1 | | 96 standard plate | |
|---|---|---|---|
| Name | Well volume [µl] | Total reagent volume [µl] | Type |
| 0.5% TFA | 100 | - | Reagent |

| TipPlate and Beads | | 96 standard plate | |
|---|---|---|---|
| Name | Well volume [µl] | Total reagent volume [µl] | Type |
| Beads | 150 | - | Reagent |

| Elution2 | | 96 standard plate | |
|---|---|---|---|
| Name | Well volume [µl] | Total reagent volume [µl] | Type |
| 1% TFA, 60% ACN | 150 | - | Reagent |

#### Steps data

| | | | |
|---|---|---|---|
| Tip1 | | 96 DW tip comb | |
| Pick-Up | | TipPlate and Beads | |
| CollectBeads1 | | TipPlate and Beads | |
| | | Collect count | 5 |
| | | Collect time [s] | 10 |
| Bind | | Peptides | |
| | Beginning of step | Precollect | No |
| | | Release beads | Yes |
| | Mixing / heating: | Shake 1 time, speed | 00:00:10, Medium |
| | | Shake 2 time, speed | 00:09:50, Slow |
| | | Loop count | 12 |
| | | Heating during mixing | No |
| | End of step | Postmix | No |
| | | Collect count | 5 |
| | | Collect time [s] | 10 |
| Wash1 | | Wash1 | |
| | Beginning of step | Precollect | No |
| | | Release time, speed | 00:01:00, Slow |
| | | Mixing | [none] |
| | | Heating during mixing | No |
| | End of step | Postmix | No |
| | | Collect count | 5 |
| | | Collect time [s] | 10 |
| Wash2 | | Wash2 | |
| | Beginning of step | Precollect | No |
| | | Release time, speed | 00:01:00, Slow |
| | | Mixing | [none] |
| | | Heating during mixing | No |
| | End of step | Postmix | No |
| | | Collect count | 5 |
| | | Collect time [s] | 10 |
| Wash3 | | Wash3 | |
| | Beginning of step | Precollect | No |
| | | Release time, speed | 00:01:00, Slow |
| | | Mixing | [none] |
| | | Heating during mixing | No |
| | End of step | Postmix | No |
| | | Collect count | 5 |
| | | Collect time [s] | 10 |

Protocol report
pY_SuperBinder_v2
9/8/2022 1:52:25 PM-07:00 4/4

| Step | | Parameter | Value |
|---|---|---|---|
| H2O Wash | | H20 Wash | |
| | Beginning of step | Precollect | No |
| | | Release time, speed | 00:01:00, Slow |
| | | Mixing | [none] |
| | | Heating during mixing | No |
| | End of step | Postmix | No |
| | | Collect count | 5 |
| | | Collect time [s] | 10 |
| Elution1 | | Elution 1 | |
| | Beginning of step | Precollect | No |
| | | Release time, speed | 00:00:30, Slow |
| | Mixing / heating: | Shake 1 time, speed | 00:00:10, Medium |
| | | Shake 2 time, speed | 00:00:50, Slow |
| | | Loop count | 10 |
| | | Heating during mixing | No |
| | End of step | Postmix | No |
| | | Collect beads | No |
| Pause1 | | Elution2 | |
| | | Message | Load Elution 2 |
| Collect From E1 | | Elution 1 | |
| | | Collect count | 5 |
| | | Collect time [s] | 10 |
| Elution2 | | Elution2 | |
| | Beginning of step | Precollect | No |
| | | Release time, speed | 00:00:30, Slow |
| | Mixing / heating: | Shake 1 time, speed | 00:00:10, Medium |
| | | Shake 2 time, speed | 00:00:50, Slow |
| | | Loop count | 10 |
| | | Heating during mixing | No |
| | End of step | Postmix | No |
| | | Collect count | 5 |
| | | Collect time [s] | 10 |
| Dispose Beads | | H20 Wash | |
| | | Release time, speed | 00:00:10, Fast |
| Leave | | TipPlate and Beads | |

### 5. R2-P2 (for clean-up of R2-pY eluate)

#### a. Reagent and Material List

| Item | Vendor | Catalog # |
|---|---|---|
| KingFisher™ Flex | Thermo Fisher Scientific | |
| KingFisher 96 KF microplate (200 µL) | Thermo Fisher Scientific | 97002540 |

| | | |
|---|---|---|
| KingFisher 96 microtiter DW plate | Thermo Fisher Scientific | 95040450 |
| KingFisher 96 tip comb for DW magnets | Thermo Fisher Scientific | 97002534 |
| SpeedVac Vacuum Concentrator | various | |
| Plate, PCR or microtube centrifuge | various | |
| Bath sonicator | various | |
| Fe-NTA MagBeads | Cube Biotech | 31501-Fe |
| Water LC-MS grade | Fisher Scientific | 7732-18-5 |
| Acetonitrile LC-MS grade | Fisher Scientific | 75-05-8 |
| Formic acid LC-MS grade | Fisher Scientific | A117-50 |
| Trifluoroacetic acid LC-MS grade | Fisher Scientific | A116-50 |
| Ammonia (ammonium hydroxide), $NH_4OH$ | Sigma-Aldrich | 221228-A |
| Ferric chloride, $FeCl_3$ (optional) | Sigma-Aldrich | F2877 |
| EDTA (optional) | Sigma-Aldrich | EDS |
| Methanol LC-MS grade (optional) | Fisher Scientific | A456-4 |
| Acetic acid glacial HPLC grade (optional) | Fisher Scientific | A35-500 |
| C8 Extraction disks 3M$^{TM}$ Empore$^{TM}$ (optional) | Fisher Scientific | 14-386 |

#### b. Protocol

##### i. Important Notes

- Any precipitate present in the peptide sample significantly reduces the selectivity of phosphopeptide enrichment and needs to be removed by centrifugation prior to the binding step.
- Do not leave magnetic beads without liquid for longer than 1 min.
- Do not leave magnetic beads in aqueous solution for longer than 2 hours.
- If magnetic beads are clumped or aggregated, sonicate them briefly in a bath sonicator.
- Magnetic $Fe^{3+}$-IMAC beads can be reused by stripping the ion metal and reloading it. We have observed no change in performance after reuse for up to 2 years.
- All plate and pipetting steps are performed with 8- or 12- multi-channel pipettes.

##### ii. Experiment Planning

- We recommend using 100 µL of 5% $Fe^{3+}$ beads in 80% ACN, 0.1% TFA for 0.25 mg to 4 mg of input sample. However, the amount of $Fe^{3+}$ beads may be varied for particularly low or high pY content samples.

- Determine the volumes of solutions to be used per well in the different plates:
  - Binding plate: 150 µL 80% ACN, 0.1% TFA per well.
  - Wash plates (3x): 150 µL 80% ACN, 0.1% TFA per well.
  - Elution plate: 50 µL 50% ACN, 2.5% $NH_4OH$ per well.
  - Acidification solution for elution plate: 30 µL 75% ACN, 10% formic acid per well.

##### iii. Magnetic Fe3+-IMAC Bead Preparation and Recycling

Magnetic bead preparation steps are performed on a magnetic rack in 2ml tubes. For washing, remove tubes from the rack, homogenize beads by flicking the tube, and remove all liquid.

- Usage of new $Fe^{3+}$-NTA MagBeads:
  - PureCube Fe-NTA MagBeads are delivered as a 25% suspension and are ready to use for phosphopeptide enrichment.
  - Before use, dilute beads to 5% and wash three times with 80% ACN, 0.1% TFA. For all further handling steps, beads are kept at 1 mL aliquots at a working concentration of 5%.
- Storage of used $Fe^{3+}$-NTA MagBeads:
  - After R2-P2, beads can be recollected before they dry out.
  - Wash once with 1 mL 50% ACN, 50% MeOH, 0.01% acetic acid.
  - Store beads in 1 mL of the same buffer at 4°C until regeneration.
- Stripping and reloading of $Fe^{3+}$-NTA MagBeads:
  - Wash beads three times with 1 mL of water.
  - Wash once with 1 ml 40-100 mM EDTA, pH 8.
  - Resuspend beads in 1 ml 40-100 mM EDTA, pH 8 and incubate for 30 minutes while shaking or rotating the tubes. Ensure that the beads remain in solution.
  - Wash beads three times with 1 ml of water.
  - Wash once with 1 ml 10 mM $FeCl_3$.
  - Resuspend in 1 ml 10 mM $FeCl_3$ and incubate for 30 min, while shaking or rotating in tubes. Ensure that the beads remain in solution.
  - Wash beads three times with 1 ml of water.
  - Wash beads three times with 1 ml 80% ACN, 0.1% TFA.
  - Resuspend in 1 ml 80% ACN, 0.1% TFA.
  - Beads are ready to use.

##### iv. $Fe^{3+}$-IMAC plate preparation for R2-P2

- Retrieve peptides from R2-pY that were brought to 80% ACN and centrifuged to remove any precipitate. If peptides were dried by vacuum centrifugation, resuspend peptides with 900 µL 80% ACN, 0.1% TFA by shaking and incubating in a bath sonicator for 10 min. Make sure that peptides go completely into solution. Insoluble precipitates need to be removed by centrifugation.
- **Tip plate:** Prepare an empty tip plate with a comb.

- **Bead plate:** Prepare one plate with the calculated amount of IMAC beads. 100 µL of beads is recommended. However, if the volume is less than 50 µL, fill up to 50 µL with 80% ACN, 0.1% TFA.
- **Wash plates:** Prepare three wash plates with 150 µL 80% ACN, 0.1% TFA per well.
- **Run program**: Start the R2-P2 KingFisher program and follow the instructions on the robot. Leave the elution plate position empty and start the protocol. The elution plate is prepared and loaded later in order to prevent evaporation of the $NH_4OH$, which could compromise elution efficiency.
- **Elution plate**: The R2-P2 Kingfisher program will pause after approximately 35 minutes. At this point pipette 50 µL of 50% ACN, 2.5% $NH_4OH$ into each well of the elution plate.
- Follow robot instructions, load the elution plate and resume the program.
- Remove all plates after the program finishes.
- Immediately neutralize the elution by adding 30 µL of 75% ACN, 10% formic acid to the plate. See optional filter step or transfer to a MS vial or MS sample plate.
- **Optional filter step:** Filter samples through C8 stage tip or stage tip plate. For this, pack a stage tip with one layer of C8 material and perform the following steps:

We recommend filtering samples when processing many samples in parallel to avoid any carryover of magnetic beads to the LC system.

  - Condition with 30 µL MeOH
  - Wash with 30 µL ACN
  - Wash with 30 µL 75% ACN, 0.25% acetic acid.
  - Position stage tip above MS sample vial.
  - Adjust sample to 50% ACN and pass sample through stage tip, collecting eluate in the MS vial.
    - Consistently, 10-20 µL evaporation from sample occurs and can be replaced by addition of 10 - 20 µL 100% ACN prior to loading sample.
  - Elute with 30 µL 70% ACN, 0.25% acetic acid, collecting this second eluate also in the MS vial.
- Dry down samples in a vacuum centrifuge. Dried samples can be stored at -20°C. Resuspend in 4% formic acid and 3% ACN prior to LC-MS analysis.

#### v. R2-P2 KingFisher™ Flex Protocol



#### General info

Protocol information

| | |
|---|---|
| Protocol name | R2-P2_FeIMAC_post-SB-enrichment_v1 |
| Modified by | KingFisher |
| Kit name | R2-P2_Fe-IMAC |
| Description | R2-P2 Fe3+ IMAC protocol for clean-up of tyrosine phosphopeptides after R2-pY.<br>[Alexis Chang, Villen Laboratory, University of Washington] |



#### Reagent info

| Beads | | 96 standard plate | |
|---|---|---|---|
| Name | Well volume [µl] | Total reagent volume [µl] | Type |
| Fe-IMAC beads in 80%ACN, 0.1%TFA | 100 | - | Reagent |

| Peptides | | 96 DW plate | |
|---|---|---|---|
| Name | Well volume [µl] | Total reagent volume [µl] | Type |
| Sample (peptides) in 80%ACN, 0.1%TFA | 900 | - | Sample |

| Wash 1 | | 96 standard plate | |
|---|---|---|---|
| Name | Well volume [µl] | Total reagent volume [µl] | Type |
| Binding buffer - 80%ACN, 0.1%TFA | 150 | - | Reagent |

| Wash 2 | | 96 standard plate | |
|---|---|---|---|
| Name | Well volume [µl] | Total reagent volume [µl] | Type |
| Binding buffer - 80%ACN, 0.1%TFA | 150 | - | Reagent |

| Wash 3 | | 96 standard plate | |
|---|---|---|---|
| Name | Well volume [µl] | Total reagent volume [µl] | Type |
| Binding buffer - 80%ACN, 0.1%TFA | 150 | - | Reagent |

| Elution | | 96 standard plate | |
|---|---|---|---|
| Name | Well volume [µl] | Total reagent volume [µl] | Type |
| Elution buffer 50%ACN, 50% 1:20 ammonia | 50 | - | Reagent |

| Tip Plate | | 96 standard plate | |
|---|---|---|---|
| Name | Well volume [µl] | Total reagent volume [µl] | Type |
| - | - | - | - |

#### Steps data

| | | |
|---|---|---|
| **Tip1** | **96 DW tip comb** | |
| **Pick-Up** | **Tip Plate** | |
| **Mix and Collect Beads** | **Beads** | |
| Beginning of step | Precollect | No |
| | Release beads | No |
| Mixing / heating: | Mixing time, speed | 00:00:15, Bottom mix |
| | Heating during mixing | No |
| End of step | Postmix | No |
| | Collect count | 5 |
| | Collect time [s] | 10 |
| **Bind** | **Peptides** | |
| Beginning of step | Precollect | No |
| | Release time, speed | 00:01:00, Medium |
| Mixing / heating: | Mixing time, speed | 00:30:00, Medium |
| | Heating during mixing | No |
| End of step | Postmix | No |
| | Collect count | 5 |
| | Collect time [s] | 10 |
| **Wash1** | **Wash 1** | |
| Beginning of step | Precollect | No |
| | Release time, speed | 00:01:00, Medium |
| Mixing / heating: | Mixing time, speed | 00:01:00, Medium |
| | Heating during mixing | No |
| End of step | Postmix | No |
| | Collect count | 5 |
| | Collect time [s] | 10 |
| **Wash2** | **Wash 2** | |
| Beginning of step | Precollect | No |
| | Release time, speed | 00:01:00, Medium |
| Mixing / heating: | Mixing time, speed | 00:01:00, Medium |
| | Heating during mixing | No |
| End of step | Postmix | No |
| | Collect count | 5 |
| | Collect time [s] | 10 |
| **Wash3** | **Wash 3** | |
| Beginning of step | Precollect | No |
| | Release time, speed | 00:01:00, Medium |
| Mixing / heating: | Mixing time, speed | 00:01:00, Medium |
| | Heating during mixing | No |
| End of step | Postmix | No |
| | Collect beads | No |

| | | | |
|---|---|---|---|
| Pause - Load Elution Plate | | Elution | |
| | | Message | Load Elution plate |
| | | Dispensing volume [µl] | 0 |
| Collect Beads | | Wash 3 | |
| | | Collect count | 5 |
| | | Collect time [s] | 10 |
| Elution of phosphopeptides | | Elution | |
| | Beginning of step | Precollect | No |
| | | Release time, speed | 00:00:30, Bottom mix |
| | Mixing / heating: | Mixing time, speed | 00:05:00, Bottom mix |
| | | Heating during mixing | No |
| | End of step | Postmix | No |
| | | Collect count | 5 |
| | | Collect time [s] | 10 |
| Dispose Beads | | Beads | |
| | | Release time, speed | 00:00:10, Fast |
| Leave | | Tip Plate | |

### Supporting Methods 2:

#### Rationale for testing different conjugation chemistries for attachment of sSrc to magnetic particles.

a. NHS chemistry targets primary amines on sSrc to form amide bonds.
b. Maleimide chemistry targets free thiols on sSrc to form thioether bonds.
c. Epoxy chemistry targets both amines and thiols on sSrc depending on solution pH. Epoxy conjugation results in secondary amine and thioether bonds, respectively. Conjugation was performed at both pHs and bead slurry was pooled in equal volumes for testing.

### Supplementary Figures

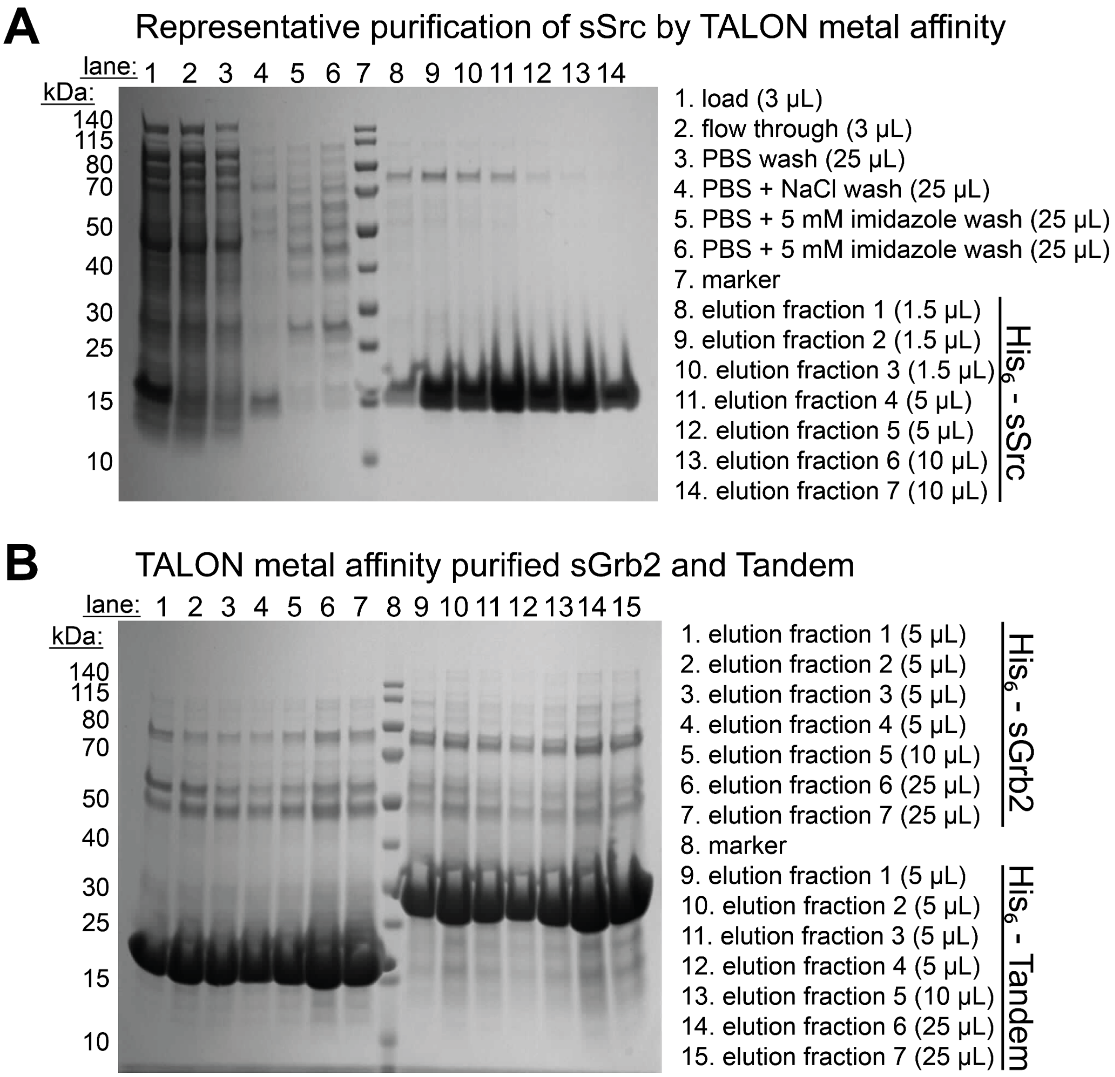


#### Supporting Figure 1. Affinity purification of pY superbinders.

**A.** Representative SDS-PAGE depicting fractions throughout purification of sSrc using TALON immobilized metal affinity chromatography. Elution fractions were pooled prior to buffer exchange and conjugation to magnetic beads (lanes 8-14). **B.** SDS-PAGE of TALON purified fractions for each sGrb2 (lanes1-7) and Tandem (lanes 9-15) pY superbinder domains. Elution fractions were pooled prior to buffer exchange and conjugation to magnetic beads.

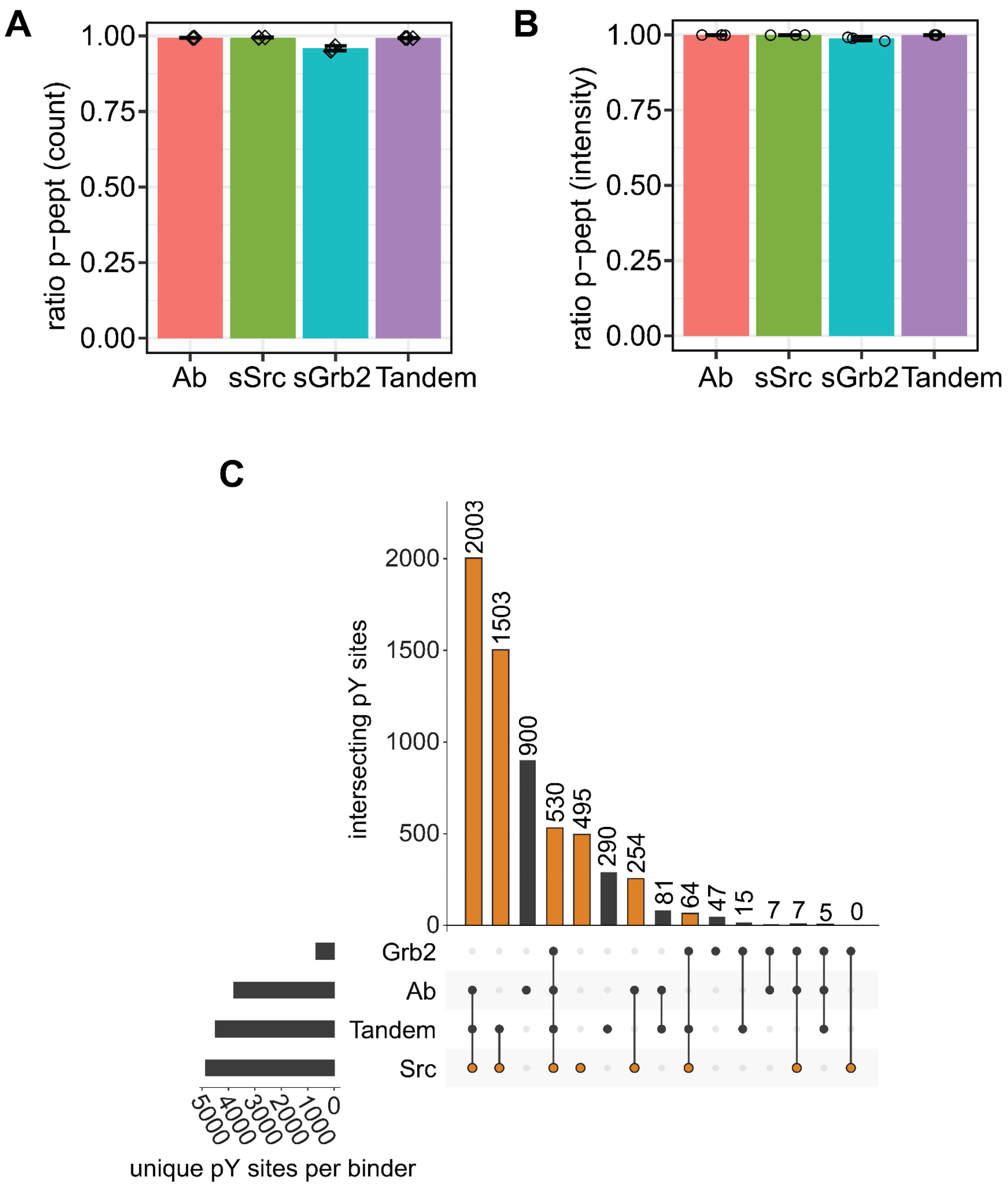


#### Supporting Figure 2. All pY binders offer high enrichment efficiency and expand the detectable pY proteome.

**A.** Phosphopeptide enrichment efficiency is shown as a ratio of the number of phosphorylated peptides over the combined count of all detected peptides. **B.** Phosphopeptide enrichment efficiency is shown as a ratio of the intensity of phosphorylated peptides over the intensity of all detected peptides (mean ± SD, *n* = 3). **C.** Barplot depicting intersections of distinct pY sites enriched by each binder after triplicate measurements. Identity of each intersection is displayed as connected dots below each bar. Total number of unique pY sites enriched by each binder are represented as a bar plot in the bottom left. Intersections representing pY sites enriched by sSrc are colored orange. pY-sites not detected following sSrc enrichment are colored black.

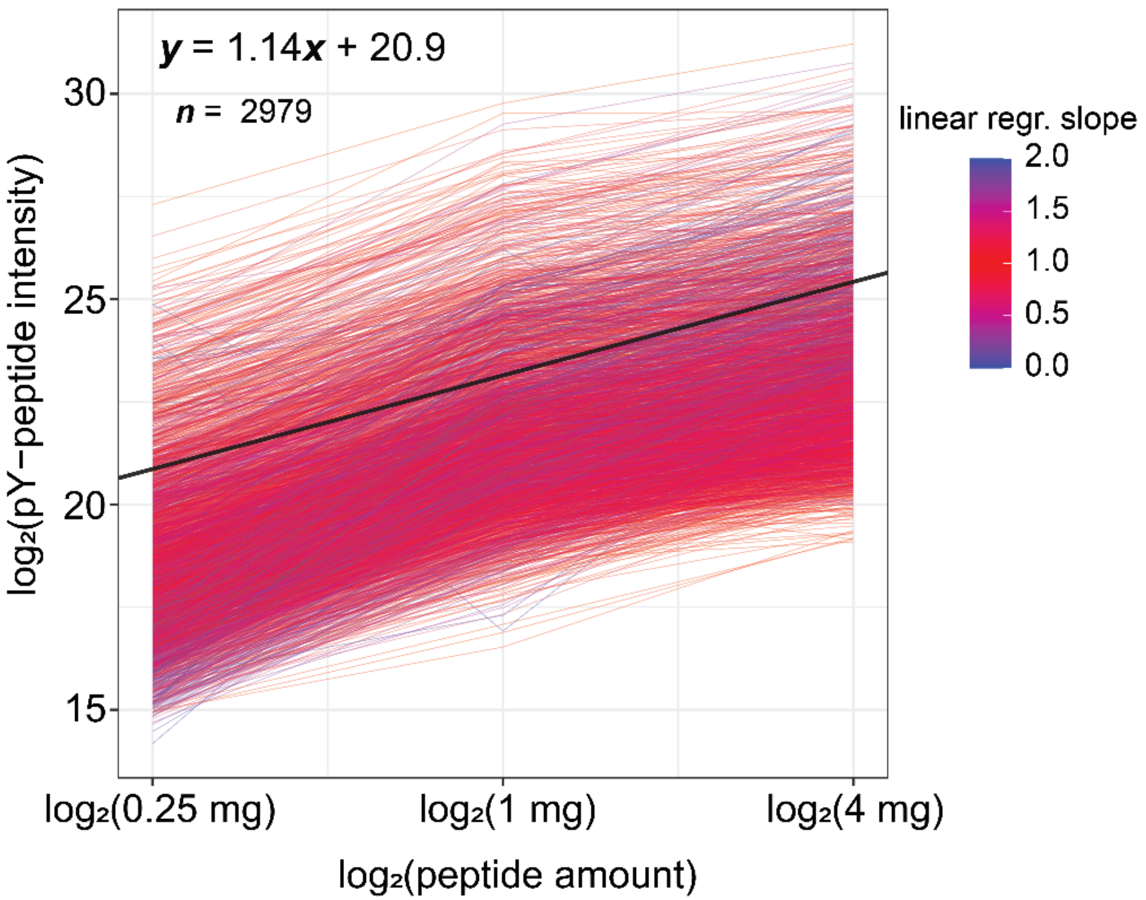


### Supporting Figure 3. R2-pY is quantitatively scalable.

Linear regressions of MS1 intensities for individual pY-peptides are plotted in pink to blue. Slopes of 1 (linear) are more red while linear regressions with slopes farther than 1 (less linear) are more blue. Black line represents the median of individual regressions. Equation of median linear regression printed in the top left of the plot.

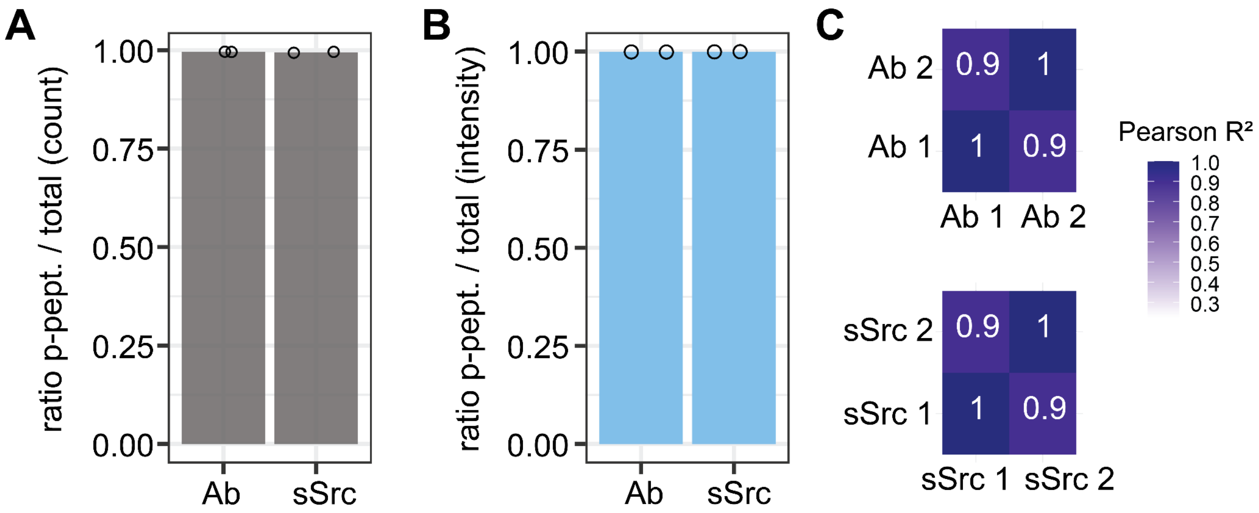


#### Supporting Figure 4. R2-P1 to R2-pY to R2-P2 displays high phosphopeptide enrichment efficiency.

**A.** Phosphopeptide enrichment efficiency is shown as a ratio of the number of phosphorylated peptides over the combined count of all detected peptides. **B.** Phosphopeptide enrichment efficiency is shown as a ratio of the intensity of phosphorylated peptides over the intensity of all detected peptides ($n$ = 2). **C.** Pearson $R^2$ correlation coefficients of pY-peptide intensity between duplicate enrichments by each binder.
